# Amplitude entropy to capture chimera-like behavior in the altered brain dynamics during seizures

**DOI:** 10.1101/2024.05.26.595969

**Authors:** Saptarshi Ghosh, Isa Dallmer-Zerbe, Barbora Rehak Buckova, Jaroslav Hlinka

## Abstract

Epilepsy is a neurological disease characterized by epileptic seizures, which commonly manifest with pronounced frequency and amplitude changes in the EEG signal. In the case of focal seizures, initially localized pathological activity spreads from a so-called “onset zone” to a wider network of brain areas. Chimeras, defined as states of simultaneously occurring coherent and incoherent dynamics in symmetrically coupled networks are increasingly invoked for characterization of seizures. In particular, chimera-like states have been observed during the transition from a normal (asynchronous) to a seizure (synchronous) network state. However, chimeras in epilepsy have only been investigated with respect to the varying phases of oscillators. We propose a novel method to capture the characteristic pronounced changes in the recorded EEG amplitude during seizures by estimating chimera-like states directly from the signals in a frequency- and time-resolved manner. We test the method on a publicly available intracranial EEG dataset of 16 patients with focal epilepsy. We show that the proposed measure, titled Amplitude Entropy, is sensitive to the altered brain dynamics during seizure, demonstrating its significant increases during seizure as compared to before and after seizure. This finding is robust across patients, their seizures, and different frequency bands. In the future, Amplitude Entropy could serve not only as a feature for seizure detection, but also help in characterizing amplitude chimeras in other networked systems with characteristic amplitude dynamics.

## 1 Introduction

Synchronization describes stable, functional relationships between temporal dynamics of correlated units [1]. This phenomenon is observed across a wide range of macro- and microscopic complex systems [2], including neuroscience, which is the primary application domain example of this work. Although correlated synchronous neural activities are crucial to cognitive brain functions [3], [4], the emergence of specific synchronization patterns in neural dynamics has also been associated with neurological disorders [5]. One prime example of such association is epilepsy, characterized by hypersynchronous neural activity during seizure events [6], [7]. Penfield et al. [8] postulated that seizures are essentially localized, highly synchronous brain activity characterized by high amplitude neural signals captured by Electroencephalographic (EEG) recordings.

Several studies corroborated that a signature of the altered brain dynamics during seizures may lie in the synchronization patterns of EEG channels[9]–[14]. Considerable research has been devoted both to understanding epileptic seizures by monitoring the brain’s electrical activities and to predicting seizures through the use of machine learning and advanced signal-processing tools, albeit with limited success [15], [16]. This implies a need to understand the mechanisms at play further while the brain is undergoing a seizure. The modern approach to understanding epilepsy stresses the importance of considering it (as well as other brain disorders [17]) as a network disease that is fundamentally driven by dynamical principles [18]–[22].

Increasingly, chimera states [23] are being used to characterize seizure dynamics. Chimera, in this context, refers to a system-state with co-existing synchronous and asynchronous dynamics, which was initially proposed by the works of Kuramoto et al. [24] and Abrams et al. [25]. Since then, the use of this term for a hybrid dynamical state has expanded across theoretical and experimental domains in the description of various complex systems [26], [27]. Relating chimera states to epilepsy, Andrzejak et al. [28] proposed a possible transitory relation between chimeric dynamics and the hyper-synchronous dynamics in neuronal states during seizure events (studying a dataset of three epileptic patients). Lainscsek et al. [29] reported a transition from a desynchronized to a highly synchronized neuronal state during seizure through a partially synchronous chimeric stage observed at seizure onset. The study further demonstrated signatures of such transitory chimera states already long before a seizure event (perhaps unfortunately for the possible clinical applications, with their specific methodology, the authors have only been able to detect chimera-like dynamics in one of all the patients in their dataset). Moreover, chimera state brain modeling could replicate seizure phenomena [30] and virtually mimic epilepsy surgery [31] by simulating the consequences of disrupting structural connectivity brain networks using empirical data of healthy subjects [32]. Such works highlight the potential of developing new and practically feasible methods to recognize chimeric states during seizure dynamics to understand, detect, predict, and treat seizures. Yet, investigations have not been conducted in large datasets of seizure recordings encompassing patients from different age groups and other clinical markers.

Furthermore, chimeric states in relation to seizures have primarily been defined with phases of neural oscillators [28]–[31] which define the synchronization (and asynchronization) in temporal dynamics of the sub-populations of the brain networks. However, estimating the instantaneous oscillatory phase in EEG experiments is a non-trivial problem. Recently, Onojima et al. [33] proposed a novel Kalman filter-based method to estimate the instantaneous phase of EEG oscillations but also reported on potential limitations of such techniques.

Here, we propose an alternative amplitude-based approach to investigate the connection between chimera-like states and the heterogeneity in amplitudes that emerge in seizure dynamics. Along with classically defined phase chimera, chimeric dynamics can manifest in amplitude variations, too, as demonstrated in several modeling studies and known as *amplitude chimeras* [34], [35]. Given the previous demonstration of (phase) chimera-like states in seizure recordings [28], [29] and the fact that focal seizures are often characterized by initially localized changes in amplitudes [36] and phase-amplitude coupling [37] in neural signals which are significantly different from the non-seizure state, we argue that there are likely amplitude chimera-like states in seizure dynamics that are detectable from EEG signals. This idea is supported by Rungratsameetaweemana et al. [38], who also reported the existence of such heterogeneity in the dynamics of brain networks prone to epilepsy.

In this work, we present a novel method aimed to capture chimera-like behavior in the analytical signal amplitudes of intracranially recorded electroencephalographic (iEEG) data. Evidence for its ability to track seizure-related dynamics is collected from 100 seizures recorded from 16 patients with focal epilepsy, which is the biggest dataset studied in the context of chimera-like states in epilepsy so far. The suggested method quantifies channel heterogeneity. It is a frequency- and time-point-resolved, entropy-based measure called *Amplitude Entropy* (AE). The AE progression is studied around the time of seizure onset and mean AE is compared between seizure intervals and before- and after-seizure intervals to demonstrate the significance of AE changes during seizures. Note, that our proposed method is not meant to be used as a standalone proof or detector of amplitude chimeras, the chimera theory rather served as an inspiration and interpretation framework in this study.

## 2 Materials and Methods

### 2.1 iEEG Data

The anonymized iEEG data used in this study is part of a public Epilepsy dataset [39], assembled at the Sleep-Wake-Epilepsy-Center (SWEC) of the University Department of Neurology at the Inselspital Bern and the Integrated Systems Laboratory of the ETH Zurich available from http://ieeg-swez.ethz.ch/. Data were recorded using strip, grid, and depth electrodes. Detailed information about the acquisition methods and patient metadata is mentioned in Ref [40], [41]. Data characteristics are listed in Table 1; more meta data is available in Supplementary Table **ST1** online. The patient-specific clinical factors, such as their diagnosis with regards to location of the epileptogenic brain area (‘type of epilepsy’, note that the table column falls short on addressing all aspects of clinical epilepsy type classification) and surgery outcome as classified by Engel score (1: excellent surgery outcome - no seizures after surgery, to 4: poor surgery outcome - seizures continue after surgery) were obtained from Burrello et al. [41].

**Table 1.** Selected patient and data characteristics. TLE Temporal Lobe Epilepsy, PLE Parietal Lobe Epilepsy. Note that patient numbering in our study deviated from the numbering in the dataset (such that for example original ID1 in Ref [40] and P1 in Ref 37 [41] became ID10).

| Patient | Age (y) | Engel outcome | No. of electrodes | No. of seizures | Seizure duration<br>Min/Max (s) | Type of epilepsy<br>& (MRI) |
| --- | --- | --- | --- | --- | --- | --- |
| ID1 | 24 | 2 | 47 | 13 | 10/252 | TLE (neg.) |
| ID2 | 19 | 1 | 42 | 4 | 96/301 | TLE (pos.) |
| ID3 | 25 | 1 | 98 | 2 | 73/125 | TLE (neg.) |
| ID4 | 32 | 4 | 62 | 14 | 31/160 | PLE (pos.) |
| ID5 | 20 | 2 | 54 | 10 | 66/154 | TLE (neg.) |
| ID6 | 48 | 1 | 64 | 4 | 89/190 | TLE (pos.) |
| ID7 | 31 | 4 | 36 | 2 | 14/16 | PLE (pos.) |
| ID8 | 38 | 4 | 59 | 2 | 52/61 | TLE (neg.) |
| ID9 | 27 | 1 | 56 | 9 | 104/198 | TLE (neg.) |
| ID10 | 46 | 2 | 100 | 5 | 10/22 | TLE (neg.) |
| ID11 | 26 | 1 | 64 | 2 | 83/135 | TLE (pos.) |
| ID12 | 59 | 4 | 49 | 10 | 23/93 | TLE (pos.) |
| ID13 | 49 | 2 | 92 | 7 | 19/100 | FLE (pos.) |
| ID14 | 36 | 1 | 74 | 7 | 154/1002 | PLE (pos.) |
| ID15 | 23 | 4 | 61 | 3 | 52/184 | TLE (neg.) |
| ID16 | 31 | 2 | 59 | 6 | 67/117 | TLE (pos.) |

The iEEG dataset includes data from 16 drug-resistant epileptic patients recorded during their pre-surgical evaluation for epilepsy brain surgery. The iEEG signals were recorded intracranially by strip, grid, and depth electrodes. Electrodes were placed for clinical reasons based on the hypothesized location of epileptogenic tissue in each individual. The data was pre-filtered in the 0.5 Hz to 150 Hz frequency band and sampled at 512 Hz. Both forward and backward filtering were used in the processing to minimize phase distortions. The dataset is prepossessed to remove the channels continuously corrupted by artifacts. It comes with annotated seizure on- and offsets obtained from visual inspection of an EEG board-certified, experienced epileptologist. Annotations of the clinically hypothesized location of epileptogenic tissue was not available. Each patient has recordings of multiple seizure events consisting of three minutes of pre-seizure segments (system state immediately before the seizure onset), the seizure segment (seizure event ranging from 10 s to 1002 s), and three minutes of post-seizure segment (i.e., system state after seizure event). All the information about the dataset is acquired from the dataset website [39] and the associated publications [40], [41].

### 2.2 Chimera states

A chimera state classically refers to the coexistence of two sub-populations of units in a network displaying disparate dynamics. Such classical chimera state was initially defined on phases (or on frequencies) [24], [25] of oscillators where, typically, one sub-population was characterized by correlated phases (phase-locked sub-population of oscillators) showing synchronous dynamics, while other population demonstrate asynchronous dynamics with un-correlated phases. Such chimera states was later expanded into multiple branches involving different dynamical aspects of various oscillator models including optical, chemical, mechanical and neuronal models. Parastesh et al. [26] presents an excellent review of different types of chimeras in great detail.

However, in this study, we explore a particular type of chimera, so called Amplitude chimera state. First introduced by Zakharova et al. [34], amplitude chimeras demonstrate chimera behaviour of having two co-existent oscillator subpopulations with disparate dynamics, shown in their amplitudes rather than their phases. Amplitude chimera state distinctively shows units where, one sub-population is oscillating with spatially coherent amplitude and the other exhibits oscillations with spatially incoherent amplitudes, i.e. the sequence of amplitudes of neighbouring oscillators are uncorrelated, in time. Such amplitude chimera state has been numerically shown for Stuart Landau Oscillator and Rayleigh Oscillator systems [42]. In the current study, epileptic seizures are considered as amplitude chimera-like state. Introducing a novel chimera-sensitive method (described in the next paragraph) we demonstrate amplitude chimera-like behavior during seizures in a large dataset of 100 seizures.

### 2.3 Analytic Amplitude (AA) and Amplitude Entropy (AE)

Figure 1 illustrates the step-by-step process for estimating Amplitude Entropy (AE) from iEEG data. The first step involves filtering the data, separately for each recording channel, using a fourth-order Butterworth band-pass filter, defined by the power frequency response; |*H* (*i ω*) |^2^ = 1/{1+ (*ω*/*ω*_*c*_) ^2*n*^}, where *n* = 4 is the filter order and *ω* is the angular frequency, and *ω*_*c*_ represents the cutoff frequencies. The applied cutoff frequencies resolve the recorded signal into specific frequency bands that are important in clinical studies. These frequency bands include: (*δ* 0.5 − 4*Hz*), (*θ* 4 − 8*Hz*), (*α* 8 − 12*Hz*), (*β* 12 − 35*Hz*), *low Ly* (35 − 80*Hz*), and *high Hγ* (80 − 150*Hz*). We note the data was pre-filtered in 0.5 150 *Hz* band in the Dataset itself.

**Figure 1.**
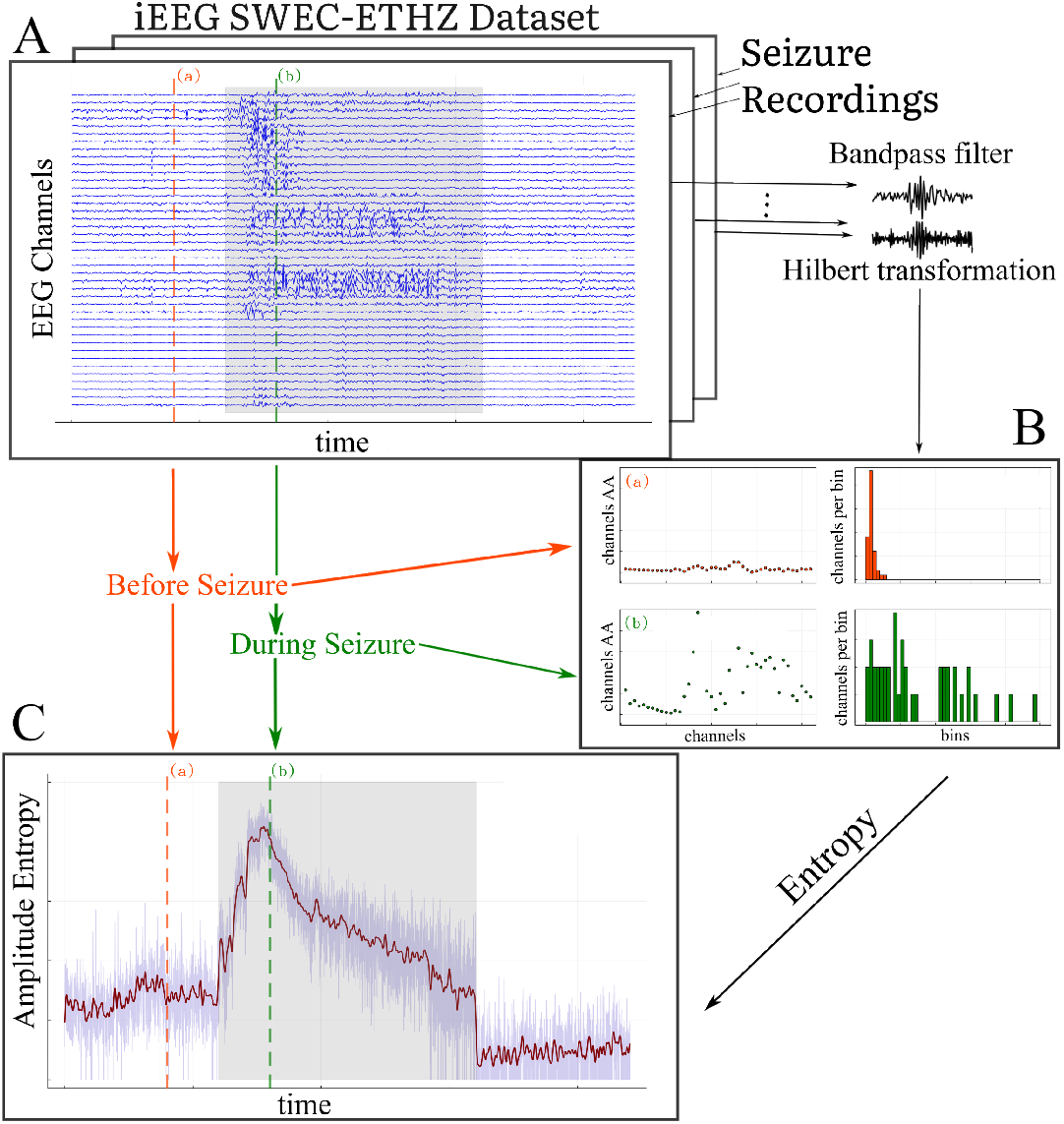
(Color Online) Workflow of computing the amplitude entropy (AE) from multi-channel iEEG data for each seizure in each patient: (**A**) provides a pictorial representation of the seizure data matrix (with time × recording-channel dimensions) taken from the SWEC-ETH database. The channels are then frequency-filtered and Hilbert-transformed to obtain the corresponding Analytic Amplitudes per frequency band (AA is the positive envelope of the waveform); (**B**) shows the distribution of AA across channels at two particular time points (time-point (a) and (b)), demonstrating an amplitude chimera-like state. This distribution of AA is tracked using entropy estimation at each time point; (**C**) displays the time series of amplitude entropy (AE) along with a time point before (marked in red) and during a seizure (marked in green). The seizure period is indicated by the gray shaded windows. The pre- and post-seizure period always spans 3 minutes (at a 512 Hz sampling rate) with a variable-length seizure ranging from 10 s to 1002 sec [41].

Next, we apply the Hilbert transformation to the filtered signal [43]. The Hilbert transform is a mathematical tool that allows us to convert the real signal into a complex-valued analytic signal, denoted as, z(*t*) = *x* (*t*) + *i*H{*x*(*t*)}, where *i* is the imaginary unit. This complex signal consists of both a real part (*x* (*t*), the original signal) and an imaginary part *i*H{*x*(*t*)}, representing a phase shifted signal. We, thereafter obtain the magnitude of the complex signal, 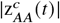 which is called the *Analytic Amplitude* (henceforth mentioned as AA) [44], for a given channel *c* of a recording. The AA represents the channel-specific instantaneous amplitude of the signal at any given moment and can be thought of as the smooth curve that follows the peaks and valleys of the (filtered)recorded signal in that channel, which we call the “envelope” of the signal. Thus, the AA is directly related to the positive envelope of the original signal, following the outline of the signal’s peaks. Mathematically, the AA is calculated as; 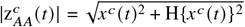, where H{*x*^*c*^ (*t*)} is the Hilbert Transform of signal *x*^*c*^ (*t*), which represents a single channel *c* of the filtered iEEG recordings. The formula essentially combines the filtered signal and its transformed counterpart to compute the instantaneous amplitude (Figure 1). The AA is particularly useful because it captures temporal variations of the amplitude of the signal for each channel of the iEEG data.

Thereafter, to quantify AA across channels at each sampled time-point, we introduce a measure called *Amplitude Entropy* (henceforth mentioned as AE) that summarizes the AA distribution across channels. Entropy, in general, is a concept that measures the degree of unpredictability or complexity in a system. Here, Amplitude Entropy helps us capture the complexity of the distribution of signal’s amplitudes and summarizes how unpredictable or diverse these amplitudes are across different iEEG channels.

To compute the entropy, at each sampled time point *t*, the AA values 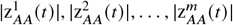 for total *m* channels (from a single seizure iEEG recording of a patient) are first binned into intervals of fixed width Δ_bin_,where the bin edges are determined by the minimum and maximum AA values across *m* channels at the sampled time-point *t* with fixed bin-width Δ_bin_. This turns the amplitude timeseries to integer timeseries of bin memberships b^1^ (*t*), b^2^ (*t*), …, b^*m*^(*t*). Let *p*_*i*_ (*t*) denote the normalized probability of the AA falling into the *i*-th bin at time *t*, estimated as the relative count:

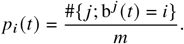

This results to in a probability mass distribution of the AA across channels of a seizure recording (Figure 1B).The entropy at time *t* is then calculated using Shannon’s entropy [45] formula:

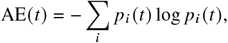

with the convention 0 log 0 = 0. The summation iterates over bins. This normalized probability distribution ensures comparability of AE values across time points, seizures, and subjects, even when the bin range (the minimum and maximum AA values across *m* channels, for a given seizure of a patient) varies due to outliers. This gives us a single, time-resolved metric that captures the complexity of the amplitude variations across channels and reflects the overall variability of the signal amplitude in each frequency band at each time-point.

We choose a binning approach with a fixed bin width Δ_bin_, which remains the same across all analysis. The bin range (min and max bin-edge) is determined by the minimum and maximum values of AA across channels at any given time-point *t*. Here we note that the bin range in our case varies but utilizing a fixed bin width ensures comparability of the entropy values across time points, seizures, and subjects (note for instance, a single outlier observation increases the bin range, but as the contribution of the empty bins is zero as 0 log 0 (in computation of entropy) is considered as 0, it does not substantially affect the entropy. For all subsequent analysis, we empirically choose a fixed bin width Δ_bin_ = 10. Other values for the bin width Δ_bin_ were tested which did not yield qualitatively different results except for scaling of the AE values (See Supplementary Figures **SF1, SF2**).

The amplitude entropy (AE) reflects the heterogeneity (or complexity) of the AA over channels for each time point. We postulate that during epileptic seizures, one may observe a separation of the AA of the iEEG channels into two sub-populations (e.g. as in Figure 1**B**) - namely channels at brain locations involved in the pathological activity, i.e. in the seizure onset zone and later the seizure propagation zone, and other channels in brain areas that are not (yet) involved - leading to an amplitude chimeric state. An amplitude chimera state characterized by co-existing subpopulations of spatially coherent and incoherent amplitude dynamics, observed during seizure can lead to long tailed AA distributions (Figure 1**B** and Figs. 2**A** and **B**) with a small number of channels with very high amplitude (belonging to spatially incoherent sub-population) and large number of channels with lower amplitudes (belonging to spatially coherent sub-population). Such amplitude chimera state state should be reflected in an increased AE in contrast with a pre-seizure state, where we expect most of the channels to exhibit AA values from much narrow distribution, leading to lower AE. Due to the individual seizure characteristics of each patient and the natural richness of recorded amplitude dynamics, however, the sub-populations are variable in time, and might be difficult to separate clearly in the AA distribution. For each patient, each seizure, and for each frequency bands, we study one time series of the AE with 3 minutes of pre and post-period and varying lengths of seizure period of the AE signal (Figure 1**C**).

**Figure 2.**
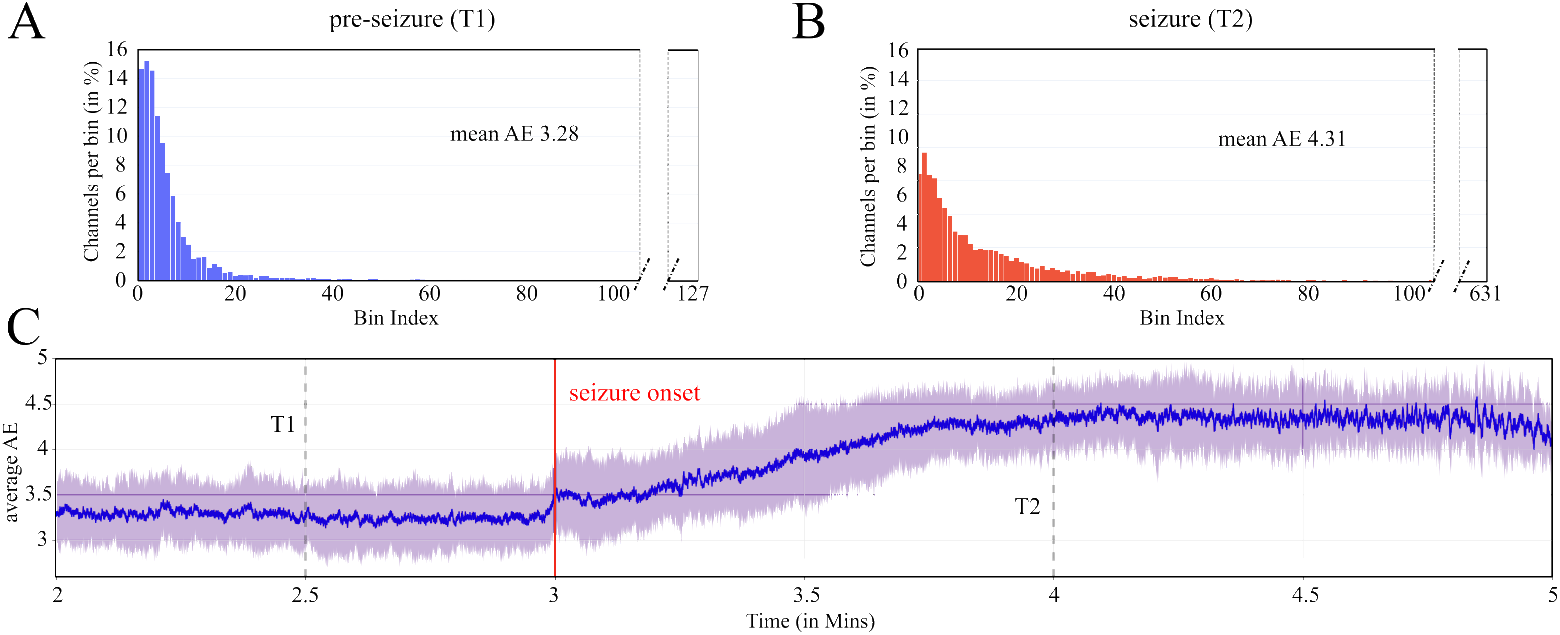
(Color Online) This figure illustrates the distribution of the Analytic Amplitude (AA) of iEEG channels (panels **A** and **B**) and the evolution of Amplitude Entropy (AE) over time (panel **C**), grand-averaged across all seizures and patients, in this order. Standard deviation averaged across patients in shades. The AA distribution exhibits a rapid decay before the seizure (**A**), in contrast to a slower decay during the seizure (**B**), indicating varying levels of heterogeneity in the AA during seizure transitions. Note also, how the total number of bins, reflecting the distance between minimum and maximum values of AA across channels, is much higher in (**B**) than in (**A**). The AE time series in (**C**) demonstrates the increase in heterogeneity during seizure transitions, characterized by the gradual decay of the AA distribution, between the periods before and during a seizure. This AE time series is computed from SWEC dataset signals which are pre-filtered between 0.5 − 150 Hz.

In order to study the time evolution of the AE (as depicted Figure 1**C** for a single seizure) averaged across seizures of different patients, the seizure segment of the AE time-series needed to be brought to equal length. We address varying seizure lengths of the seizures by adding *NaN padding* after the end of the seizure. NaN (Not a Number) typically represents missing or undefined data-points and is commonly used as a placeholder when performing numerical mathematical operations, skipping missing items. For example, in a typical averaging procedure, the divisor is adjusted at each compute-point in the presence of NaN values (i.e. with varying number of samples). Adding NaN padding to AE time-series ensured that the AE time series of all seizures were of the same time length (with NaN padding). A two-step averaging procedure (skipping NaN entries) for each time point was carried out, first across seizures and then across patients, ensuring an equally weighted contribution from all patients irrespective of their different counts of seizures. For single seizure contribution to the average per patient and then across patients, this means that, due to the NaN padding, in the beginning of the seizure interval up until shortest seizure length all seizures contribute to the average, later fewer and fewer seizures contribute until only the longest seizure contributes at the end end of the interval. We note the existence of alternative approaches in literature, e.g. as in Ref. [46] involving changing the sampling frequencies of signal segments used for length matching of seizure segments across seizures and patients. There was no noticeable difference to our results when utilizing the above-mentioned approach (see Supplementary Figures **SF3, SF4** showing time series of averaged AE when using the time normalization approach with regards to seizure duration as an alternative to the NaN padding procedure, as used in Ref. [46]).

## 3 Results

### 3.1 Amplitude entropy captures chimera-like dynamics during seizures

To study the overall, grand average AE change during the transition into seizure, we start the analysis from the broadband signal, i.e. the unmodified SWEC recordings that are pre-filtered between 0.5 − 150 *Hz*. We compute the AE at each time point with the processing pipeline described in Section 2.3 (but without the filtering stage). We then choose two time points in the time interval before the seizure (*T1*) and during the seizure (*T2*), respectively. At both of these time points, we calculate the probability mass function of the AA (pmf-AA for T1 and T2) across the channels for a given seizure and then average over all seizures and patients (see Figure 2, as well as Supplementary Figures **SF5, SF6** for patient-specific AA and AE, as well as Supplementary Figure **SF7** grand-averaged AE including also the post-seizure interval, online).

Figure 2**A** and Figure 2**B** show the averaged distribution of the AA for the two time points marked in Figure 2**C**. Comparing these two time-points, capturing the pre-seizure and seizure dynamics, aggregated over all seizure and patients, we observe a fast decaying AA distribution with a shorter tail before the seizure (Figure 2**A**), and a slow decaying AA distribution with a long tail during seizure (Figure 2**B**). This demonstrates that before the seizure most of the channels are confined within a very small bin-range showing a state similar to Figure 1**B**, top row. However, during the seizure several channels record very high amplitudes while the others record very low amplitudes (similar to Figure 1**B**, bottom row), leading to an amplitude chimera state. Such distributions also lead to lower entropy during pre-seizure periods (with lower heterogeneity across channels) and a higher entropy during seizures. Figure 2**C** shows an overall increase in the amplitude entropy while transitioning from pre-seizure to seizure state. We further observe a local peak in the AE at the seizure onset (at three min time-point of the recordings). This local peak is present in about half of the patients (see patient-specific Supplementary Figure **SF6**, online). It is due to seizure onset-specific patterns in the raw signals (see Supplementary Figures **SF8-SF13** and **SF14** for the peak for a particular case, online). We cautiously suggest that such peak could be used as a feature to detect seizure onsets, however, after taking into account the individual patient-specific variations. Later, after seizure onset and during seizure spread from the onset zone into a wider brain network, AE is increasing. Maximal AE occurs and plateaus at a time-point within minutes after seizure onset. We speculate, that the beginning of the plateau could be the time point when the maximum number of involved brain areas has been recruited.

To demonstrate the general dynamics of amplitude entropy during seizure, we define three time periods: pre-seizure, seizure and post-seizure. Note that seizure length is variable both across and within patients and that each patient has a different number of seizure recordings (Table 1). We thus compare the averaged AE (computed first over time, then over seizures, and then over patients) for the three time periods as mentioned above. Given the observed increment in AE during the seizure, we expect a triangular shape where the averaged AE during the seizure would be highest as compared to pre and post. The expected triangular shape is found in all frequencies (Figure 3), demonstrating the higher AE during seizure compared to the pre or post time periods. For patient-resolved AE points see Supplementary Figure **SF15** online.

**Figure 3.**
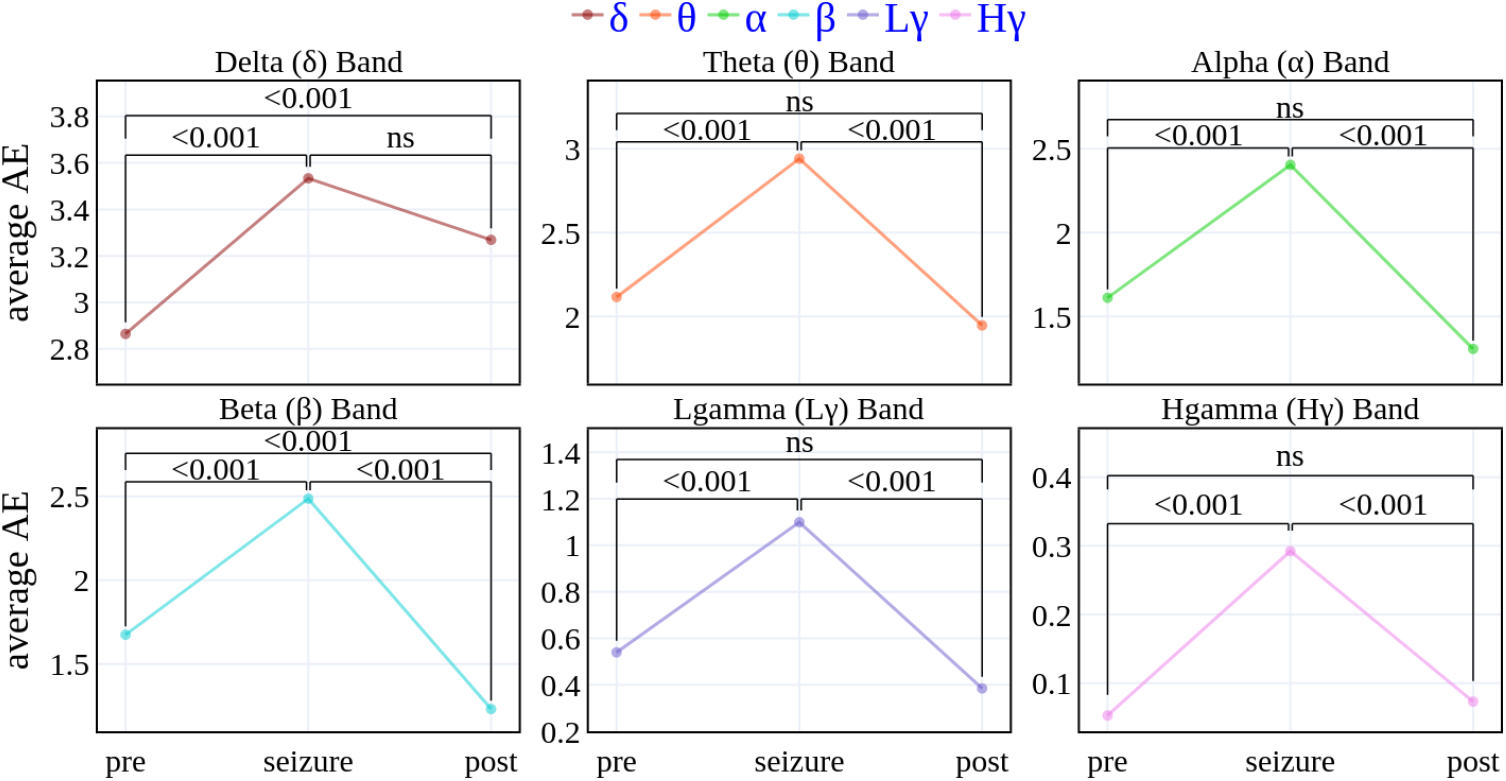
(Color Online) This figure depicts the averaged Amplitude Entropy (AE) per frequency band before, during, and after seizure. The individual AE points are first averaged over seizures, then over patients (as illustrated in Figure 1), and finally, over time windows corresponding to before, during, and after a seizure. The AE points exhibit a *statistically significant* change during seizures compared to the time periods before or after the seizure.

We evaluate the significance of AE changes across the segments (pre-seizure, seizure, post-seizure) using a linear mixed-effect model implemented in Matlab R2020b (fitlme function) [47]. The rationale for using a linear mixed-effects model lies in its ability to appropriately handle hierarchical data, such as repeated measures from the same individuals, leading to more reliable inference [48]. Average amplitude entropy served as a dependent variable and segment label (pre; seizure; post) as fixed effects. To account for variability across subjects, we included them as random effects with varying intercepts. Models were evaluated using the seizure amplitude entropy as a reference. Consequently, the null hypothesis tested was that AE during the seizure segment does not differ significantly from the pre- or post-seizure segments. The significance of the fixed effects was assessed using asymptotic Wald tests, which determine whether the estimated coefficients differ significantly from zero. Significant nonzero coefficients (below significance level of *α* = 0.05) of individual fixed-effect categories (pre vs reference, post vs reference) were considered as significant change between the reference and fixed-effect category and is annotated in Figure 3. We thus demonstrate the statistically significant change in average AE during seizure as compared to pre and post data-points.

### 3.2 Patient and frequency variations in AE seizure effects

In a post-hoc analysis, we studied the AE seizure effect as the change from pre-seizure to seizure only, calculated as difference (seizure - pre), as opposed to all three comparisons in the main results. Given the heterogeneity between patients and their seizures, that is well-documented in the literature, we were interested to see whether amplitude chimera-like states were specific to certain individuals, certain frequency bands or some clinical sub-groups of patients. For the latter, we correlated the AE seizure effect with the clinical factors available in the database. Significance testing was conducted using two-sample t-testing for the AE seizure effect between clinically different patient groups (non-parametric testing in a control analysis yielded same results), with significance level of *α* = 0.05.

Figure 4 provides a general overview using box plots for each patient and frequency band across a patient’s seizures (each dot is a seizure; please see Supplementary Figure **SF16** online, for additional separated frequency-scaled presentation). The patient-specific variations descriptively display no clear trend on a dominant frequency pattern across patients. Across frequency bands, in most patients, we observe a positive mean corroborating the findings in Figure 3.

**Figure 4.**
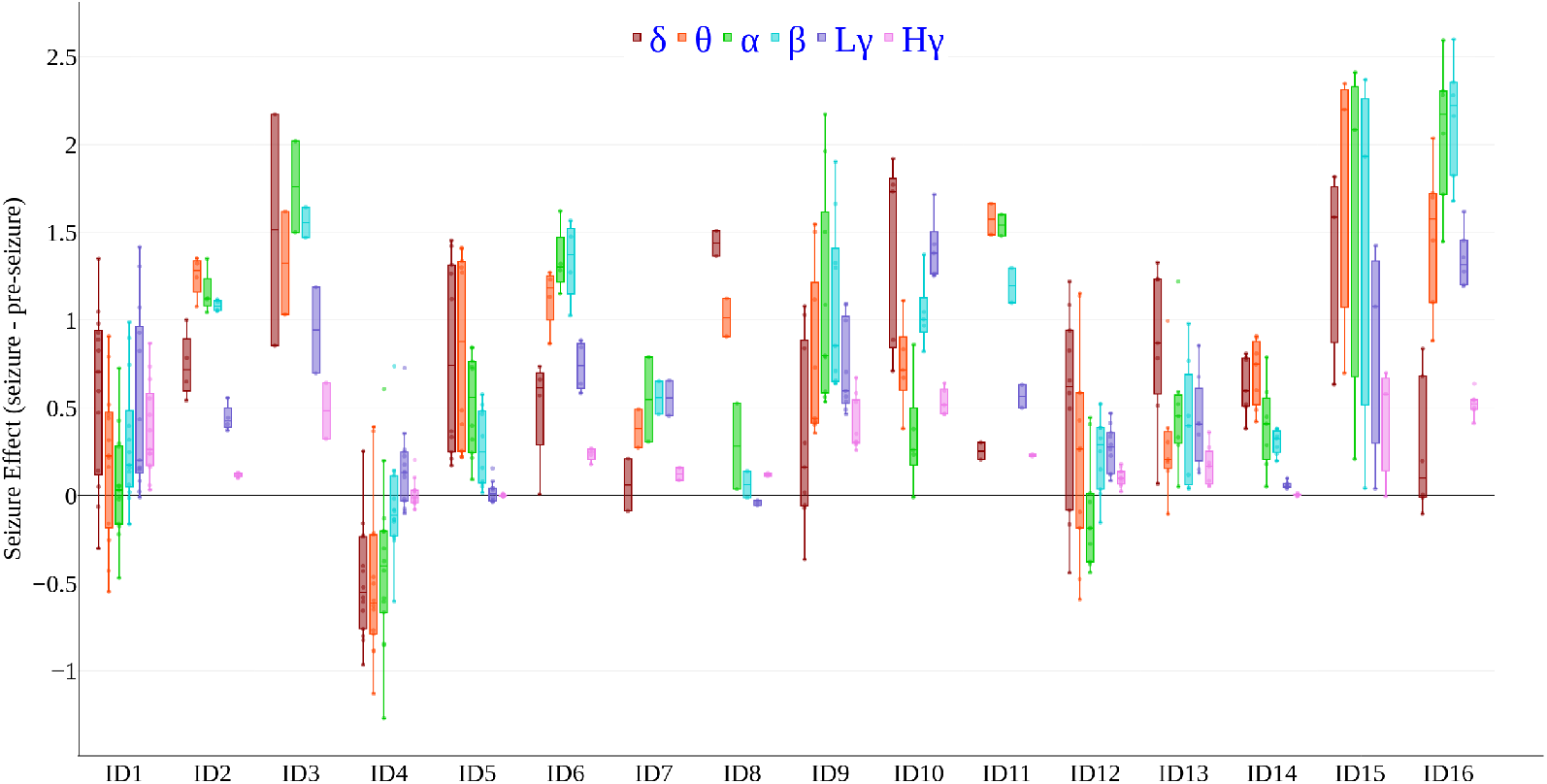
(Color Online) Seizure effect on Amplitude Entropy (AE) across patients and frequency bands. Plotted on the Y axis is the descriptive seizure effect, calculated as the difference in average AE (calculated on filtered AA of all iEEG channels per patient) between seizure and before-seizure time periods (AE averaged in time). The patient-specific seizure effect distribution is plotted for each frequency band separately (color-coded, one dot per band per seizure). The X axis represents IDs for different patients. There is an overall increase in AE from pre-seizure to seizure (values > 0), that is not frequency-specific. The frequency of maximal seizure effect is variable across patients. High gamma shows smaller variation across seizures.

Notably, clinical variables (Table 1) epilepsy type (temporal, parietal or frontal lobe epilepsy) and MRI investigation (MRI positive or MRI negative) showed a link to the AE effect during seizure (Figure 5**A**). Patients with temporal lobe epilepsy and particularly those with a positive MRI finding (typically Hippocampal Sclerosis) increased significantly more in their AE score from pre-seizure to during-seizure than patients with extratemporal seizures (*μ*_TLE_ = 0.79, *σ*_TLE_ = 0.40; *μ*_other_ = 0.24, *σ*_other_ = 0.30; two-sided Wilcoxon rank sum test with *p* < 0.05). However, given the small group sizes (12 temporal vs. 4 extratemporal epilepsy patients), a further investigation in a bigger patient sample is required to validate this finding.

**Figure 5.**
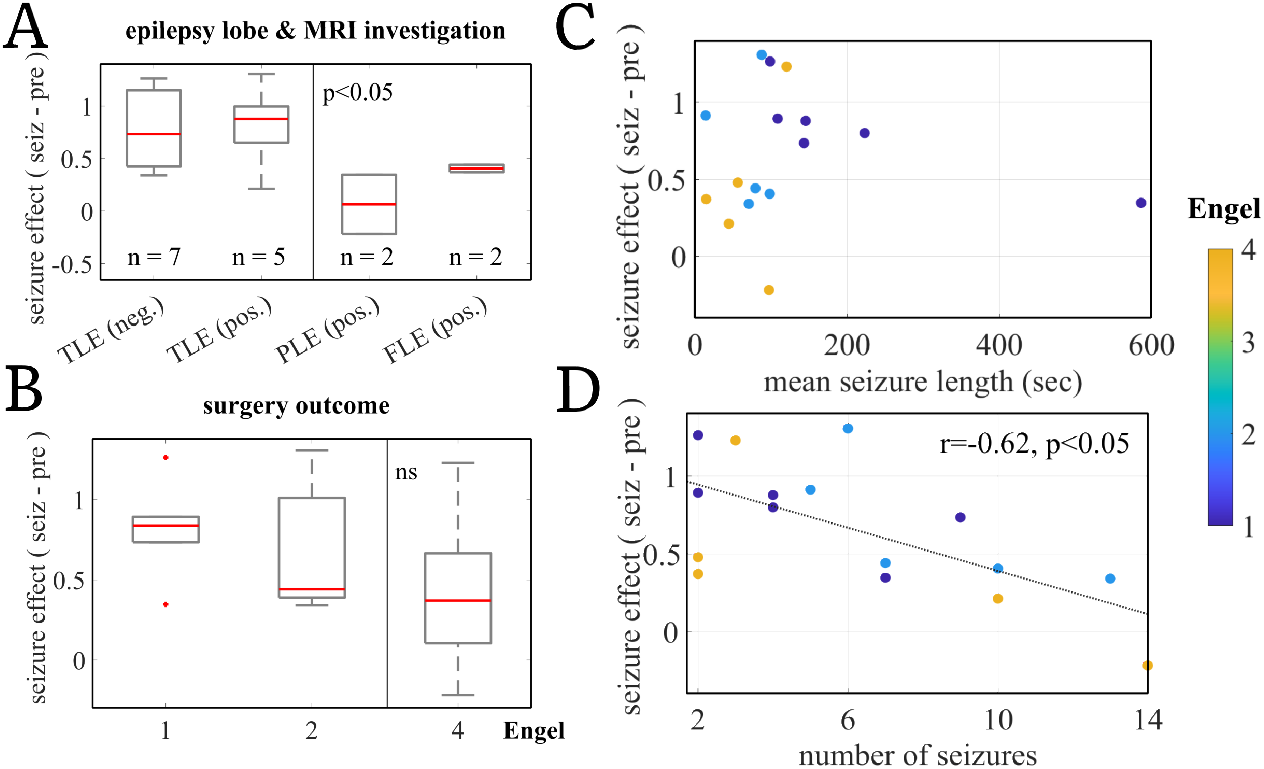
(Color Online) Seizure effect on Clinical outcomes and Seizure metadata. The seizure effect is the difference between AE points corresponding to the seizure and before-seizure time periods (averaged). (**A**) Seizure effect differs among different clinical patient groups. Patients with TLE with and without MRI finding show higher effect than extratemporal epilepsy patients. The comparison is highly influenced by patient ID4 in PLE (pos.) group. Investigation in a bigger dataset is required. (**B**) Surgery outcome shows a relationship with AE seizure effect. Patients with poor surgery outcome have lower mean AE (top). (**C**) Patients with longer mean seizure length, have better surgery outcome. (**D**) AE seizure effect negatively correlates with the numbers of seizures per patient. TLE: temporal lobe epilepsy; PLE: parietal lobe epilepsy; FLE: frontal lobe epilepsy; MRI investigation negative (neg.) means no abnormal finding on MRI. Engel scale from (1) patient is seizure free after surgery to (4) has no seizure rate improvement.

We further found a tentative relationship between the AE increase from pre-seizure to seizure and a patient’s surgery outcome (Figure 5**B**); Engel-classified surgery outcome with 1 = seizure freedom after surgery, to 4 = no improvement after surgery). Poor surgery outcome is commonly explained by an insufficient resection or a wrong clinical hypothesis on the seizure origin in the brain and thus remaining epileptogenic tissue after surgery. Insufficient resection or wrong clinical hypothesis on seizure origin location would indicate a lower accuracy of electrode placement for clinical examination of the patient, which would mean that seizure-related processes would be captured less well by the recording electrodes than in patients with good surgery outcome and thus correct clinical hypothesis on the seizure origin. In this study, patients with poor surgery outcome (n = 5) descriptively showed a lower AE increase from pre-seizure to seizure than those with an improvement after surgery. One interpretation of this would be, that, in line with the above, in those patients, the iEEG signal might have captured less of the on-going brain dynamics during seizure due to being ill-placed. However, statistical testing of a higher AE change in good vs. poor surgery outcome did not yield a significant difference (*μ*_*bad*_ = 0.41, *σ*_*bad*_ = 0.52; *μ*_*other*_= 0.76, *σ*_*other*_ = 0.34; one-sided Wilcoxon rank sum test with *p* > 0.05). Interestingly, patients with longer seizures had better surgery outcomes (*μ*_*good*_ length of seizures = 218 sec, *σ*_*good*_ = 186 sec; *μ*_*other*_ = 69 sec, *σ*_*other*_ = 36 sec; two-sided Wilcoxon rank sum test with *p* < 0.05) (Figure 5**C**). This finding could further support the outlined link between surgery outcome and AE change. Longer seizure recordings might partially be explained by a higher accuracy of electrode placement, such that seizure activity was registered in full, leading to a higher AE in cases of good surgery outcome.

Finally, we found that the AE effect was significantly correlated with the number of recorded seizures per patient (*r* = -0.62, *p* < 0.05): the more seizures were available per patient, the lower was the AE change from pre-seizure to seizure (Figure 5**D**). Previous studies have shown that patients who experience more frequent seizures may have a more complex pathology, involving diverse epileptogenic lesions and extensive seizure propagation pathways, where seizures can originate from and spread across multiple brain regions [49]. In such cases, the observed iEEG patterns might be more variable across seizures, leading to a weaker AE change from preseizure to seizure after averaging across seizures.

## 4 Discussion

This study aimed to investigate amplitude chimera-like states in epilepsy using a novel marker of chimera-like behavior that quantifies amplitude entropy across different recording sites. It shows that seizure dynamics manifest in changes of amplitude heterogeneity across the different brain areas chosen for clinical examination. Particularly, during seizure, amplitude entropy is significantly increased as compared to before and after seizure. We believe, that this finding extends the perspective of chimera-like states in epilepsy to be a phenomenon not only observed in phase synchronization of oscillators but in their amplitudes, as well. In contrast to the documented hyper-synchronization [8], [13], i.e. increased phase homogeneity in the network during seizure, we find the amplitudes in the network during seizure to be significantly more heterogeneous. This could indicate an anti-correlation of phase- and amplitude-related changes during seizure.

The finding was robust across different frequency bands. Note the difference in scaling for different frequencies, that stems from the 1/f law (also known as pink noise or flicker noise) character of EEG signal. This is a phenomenon frequently observed in recorded biological signals such as EEG where the power (and thus amplitude) decreases inversely with frequency. Thus, our use of a fixed bin-width for all the frequencies resulted in having higher counts per bin for lower frequencies compared to higher frequencies, resulting in higher relative value for AE in lower frequency signals. Therefore, bin-width is acting as a scaling factor for the computation the AE in the frequency resolved signals. However, our statistical tests prove that the seizure effect is significant also in high frequency bands, showing the robustness of the measure.

Studying the amplitude network organization across time during the transition from pre-seizure to seizure (Figure 2**C**), we find a local peak in channel heterogeneity at seizure onset (which we believe to capture some of the brain dynamics involved in seizure initiation) followed by a steady increase of AE until a plateau is reached later during seizure; in few subjects and/or seizures followed by gradual decrease. We interpret this finding as that AE is capturing the network recruitment during seizure propagation from a focal onset zone into a wider brain network. Previous studies on chimera-like states in iEEG recordings of epilepsy patients find that the outbreak of global synchronization is preceded by a transient decrease of synchronization [28]. Furthermore, Lainscsek et al. [29] report a network reorganization around seizure onset, in which a pattern of asynchronous focal onset channels and synchronous non-onset channels before seizure flips when the non-onset channels become increasingly asynchronous during seizure. Similarly, Espinoso et al. [50] show how, during seizures, focal onset channels increase while non-onset channels decrease their large-scale phase-locking. In the current study, we find a chimera-like network reorganization during seizure that manifests in an increased amplitude heterogeneity during seizure, extending the previously identified network changes to not only be a matter of phases but also of amplitudes. Future work could aim to link these findings to other evidenced changes during the transition to seizure[38], [51]–[53], including changes in phase synchronization [9], [11], [50], and in particular to studies investigating the same dataset as this study [50], [54], [55]. Furthermore, it would be interesting to study a possible relation between the chimera-like behavior and the slow dynamics that govern fluctuations in seizure likelihood over time [52], [53], [56]. A limiting factor in interpreting some of the amplitude heterogeneity across channels was the unavailability of information on which of the recording channels belongs to which electrode type (strip, grid, or depth electrodes). It is known that different types of electrodes record different types of signals [57], but also share similar signal characteristics [58]. However, as the current study focuses on dynamics, i.e. comparing time points (e.g. pre-seizure to seizure) that are equally affected by the recording differences, we do not consider this limitation critical to the validity of our main findings. Furthermore, the dataset does not provide seizure onset zone labeling, which would be valuable to compare whether a similar pattern as that suggested by as well as Espinoso et al. [50] or Lainscsek et al. [29] (asynchronous focal onset channels, synchronous non-onset channels before seizure, then non-onset channels becoming increasingly asynchronous during seizure) could be found regarding the amplitude-related network changes in this study.

While previous studies investigating chimera-like dynamics in epilepsy report their findings in few patients only (one patient in the case of Lainscsek et al., selected among a total of 15), the amplitude phenomenon as studied here was robust across patients and their seizures. Importantly, however, also this study’s findings varied on the individual patient level, such that, e.g., individual AE progressions did not always show the local peak and consecutive increase at seizure onset. Interestingly, we were able to link the size of increase from pre-seizure to seizure AE to a individual’s patient and data characteristics. Our findings suggest that chimera-like states in epilepsy could be more likely to occur in temporal lobe epilepsy patients, and especially those with a positive MRI finding such as hippocampal sclerosis. Furthermore, patients with better surgery outcome, and therefore likely a better electrode placement to help distinguish their seizure onset zone, generally showed a higher AE seizure effect. These findings, linking clinical variables to changes in iEEG features, are in line with previous studies analyzing the same database [54], [55]. Thus, they can be taken as a good sign that AE is in fact capable of capturing seizure-related brain dynamics. However, bigger patient samples, and thorough feature comparison are needed to extend these preliminary findings.

### 4.1 Amplitude chimera-like states in epilepsy

In this article, we take a complementary approach to study chimera states in epileptic seizures. We report an amplitude chimera state during seizures where we show an increase in heterogeneity (and thus amplitude entropy) in values across iEEG channels. Similar observation, correlating amplitude variations in unihemispheric sleep [59] and chimera states most famously is reported by Rattenborg et al. [60]. However, in a recently published critical review, Haugland [61] pointed out the broadening of the definition of chimera states and its increasingly weaker connection to the ‘classical’ model-based chimera states, which (as mentioned in the review) especially holds true for *natural-world* chimeras i.e. the observation of chimeric states in beyond-laboratory environments. Haugland further includes own attempts to relate chimera to unihemispheric sleep [62] such as other works like Ramlow et al. [63] reporting chimeras in relation to brain-inspired networks and dynamics [64]–[66]. Similar concern to the one of Haugland was voiced by Omelćhenko in another older review declaring all such observations ‘rather speculative’ [67].

In the context above, we would like to state our agreement with the perspective and comment that the reported amplitude chimera state in our study is rather difficult to link to model-based reports of the amplitude chimera states. We have presented an observation where spatially localized high-amplitude seizure event lead to a heterogeneous distribution in AA which can be interpreted as two (or more) sub-populations having disparate dynamics.

### 4.2 Potential applications of AE

AE changes showed a local peak at seizure onset and the seizure-related AE changes identified in this study were stronger in patients with temporal lobe epilepsy and those with better surgery outcome. These results provide first indication that AE estimation might be of clinical relevance to help not only seizure onset detection, but even patient classification. Furthermore, other network disorders bearing such variations in recorded bio-signal could benefit from the amplitude entropy approach.

It is well documented that epilepsy patients and their seizures display high variability. Therefore, there have been on-going efforts to classify epilepsy patients and their seizures into more consistent types [68], [69]. In this study, patients with temporal lobe epilepsy (with negative and positive MRI finding) showed a stronger modulation of AE than extra-temporal epilepsy patients. This opens the avenue for a potential application of AE assessment for patient classification to be investigated in follow-up studies with larger sample sizes. However, this study did not account for variability in AE due to more detailed patient and seizure characteristics, even though, for example, single seizure data visually shows a variability among the observed seizure patterns (see Supplementary Figures **SF8-SF13** online). While the goal of this study was to demonstrate the general applicability of AE assessment to epilepsy phenomena across patients and seizure types, further studies could investigate AE dynamics with regards to different types of seizures and its potential to help clinical onset marking by automatized seizure characterization.

Notwithstanding a multitude of modeling studies in chimera identification using EEG phases, we argue that real-time estimation of the instantaneous phases across channels in EEG recording is a non-trivial technologically complex problem. There have been some approaches that indeed demonstrated real-time estimation of EEG phases [33]. But such approaches are limited, relatively technologically expensive and far from being implementable in a closed-loop system. AE provides a simpler, straightforward measure to capture the amplitudes variations the emerge during seizures. Furthermore, EEG is a multi-cellular recording approach that might capture a population of neurons with varying degrees of signal-to-noise ratio. The instantaneous estimation of phases for neural oscillations may be erroneous where the baseline signal is noisy and non-oscillatory. This may make such phase-based approaches not feasible for developing seizure monitoring systems. We argue that instantaneous amplitude is a direct and far more feasible measure of the on-going brain dynamics during seizure that are captured by EEG recordings. Despite the noise in the signals, the large amplitude-heterogeneous nature of the seizure events makes it a more enticing fundamental feature for seizure characterization.

Finally, focal seizures are primarily localized in a brain area, clinically known as the epileptogenic zone. Localizing this zone is another non-trivial problem and an active area of research [70]. Electrode placement of the EEG setup in a patient may not always overlap with the epileptogenic zone, resulting in a recording where some electrodes may capture high amplitude synchronous activity from seizure areas and others might record from background asynchronous activities from other parts of the brain. Thus, our approach on one hand presents a far more feasible method for the identification of seizures, while on other hand it could potentially help the identification of epileptogenic zones from the tails of the AA distribution as shown in Figure 2. While in this study a comparison of the AA-predicted epileptogenic zones against clinically marked epileptogenic zones was not possible due to missing annotations in the public dataset, follow-up investigations should target the potential usefulness of AE and/or AA assessment in the presurgical evaluation of epilepsy patients.

## 5 Conclusion

This study provides a novel entropy-based measure to capture the emergent amplitude chimera states and conducts a data-driven investigation into the high amplitude seizure events as possible description of the brain dynamics around seizure occurrence. We show that our measure is robust and statistically significantly increased during the seizure periods in 100 seizures from the 16 patients. We thus support and extend the literature on amplitude chimera-like states in epilepsy and provide an efficient method for its detection in real iEEG data, which we hope could bear a potential to help seizure or patient classification in the future.

## 6 Data availability statement

The dataset analyzed during the current study is a publicly available dataset at http://ieeg-swez.ethz.ch/.

## 7 Code availability statement

Data processing and analysis were conducted in Python *3*.*x* and Matlab *R2020b* and *R2021a*. Relevant code and dependencies are publicly available at the COBRA group GitHub repository at https://github.com/cobragroup/amplitude-chimera-epilepsy.

## 8 Acknowledgements

This work was supported by the Czech Science Foundation (project No. 21-32608S), the Charles University Grant Agency (80120), the long-term strategic development financing of the Institute of Computer Science (RVO:67985807) of the Czech Academy of Sciences, and by ERDF-Project Brain dynamics, No. CZ.02.01.01/00/22_008/0004643.

S.G. thanks Yasser Roudi, Kavli Institute for Systems Neuroscience - NTNU, Giacomo Indiveri, Institute of Neuroinformatics - UZH and Institute of Computer Science of the Czech Academy of Sciences for support during this research collaboration.

## 9 Author contributions statement

S.G., and J.H. conceived the study and developed the methodology. S.G. and I.D.Z analyzed the data and wrote the manuscript under the supervision of J.H. B.R.B conducted the statistical analysis and reviewed the manuscript.

## 10 Additional Information

### 10.1 Competing interests

The authors declare no competing interests.

## Notes

https://github.com/cobragroup/amplitude-chimera-epilepsy

### Competing Interest Statement

The authors have declared no competing interest.

### Summary of Updates

Supplementary Figure numbers are updated to maintain sequential order

