## Supplementary material for "Amplitude entropy to capture chimera-like behavior in the altered brain dynamics during seizures": Supplimentary File

### This PDF file includes:

Table ST1 Patient Characteristics Table

Figs. SF1 Time interval- averaged Amplitude Entropy (AE) for pre-seizure, seizure and post-seizure intervals across patients per frequency band for differently chosen bin size.

Figs. SF2 Time-evolution of averaged AE (grand-averaged over seizures and patients) for differently chosen bin size.

Figs. SF3 Time-evolution of averaged AE (grand-averaged over seizures and patients) [0 1 Normalization]

Figs. SF4 Time-evolution of Amplitude Entropy (AE) per patient; AE time-series are averaged over seizures for each patient. [0 1 Normalization]

Figs. SF5 Distribution of the Analytic Amplitude (AA) of iEEG channels at T1 & T2 time-points as marked in Fig. SF6, averaged across seizures for each patients.

Figs. SF6 Time-evolution of Amplitude Entropy (AE) per patient; AE time-series are averaged over seizures for each patient. [with NaN padding]

Figs. SF7 Time-evolution of averaged AE (grand-averaged over seizures and patients) [with NaN padding]

Figs. SF8-SF13 iEEG signals for **first and second seizures** of all patients.

Figs. SF14 Local peaks in analytic amplitudes for different channels for Patient 2, Seizure 3.

Figs. SF15 Frequency-resolved averaged Amplitude Entropy (over seizures) before, during, and after a seizure with a patient-resolved and frequency-resolved grid.

Figs. SF16 Frequency-resolved and patient-resolved seizure effect on Amplitude Entropy (AE) across patients presented in frequency separated and non-scaled fashion. The seizure effect is the difference between AE points corresponding to the seizure and before-seizure time periods (averaged).

Supplementary Table ST1: Clinical factors of the Seizure Patients in SWEC dataset as collected from Burrello et al. 2020 (DOI:10.1109/TBME.2019.2919137) and renamed to match SWEC dataset IDs. Note that patient numbering in our study deviated from the numbering in the dataset (such that for example original ID1 in Burrello et al. 2018 and P1 in Burrello et al. 2020 became ID10 in our study.)

| Patient ID | Electrodes | Seizures | Duration (in s) | | Age | MRI Inves. | Epilepsy Hem.- and Lobe |
|---|---|---|---|---|---|---|---|
| | | | Min | Max | | | |
| ID1 | 47 | 13 | 10 | 252 | 24 | n | Left-Temporal |
| ID2 | 42 | 4 | 96 | 301 | 19 | y (Hippocampal Sclerosis) | Left-Temporal |
| ID3 | 98 | 2 | 73 | 125 | 25 | n | Right-Temporal |
| ID4 | 62 | 14 | 31 | 160 | 32 | y (focal cortical dysplaisa) | Left-Parietal |
| ID5 | 54 | 10 | 66 | 154 | 20 | n | Right-Temporal |
| ID6 | 64 | 4 | 89 | 190 | 48 | y (Hippocampal Sclerosis) | Left-Temporal |
| ID7 | 36 | 2 | 14 | 16 | 31 | y (tuberous sclerosis) | Right-Frontal |
| ID8 | 59 | 2 | 52 | 61 | 38 | n | Left-Temporal |
| ID9 | 56 | 9 | 104 | 198 | 27 | n | Left-Temporal |
| ID10 | 100 | 5 | 10 | 22 | 46 | n | Right -Temporal |
| ID11 | 64 | 2 | 83 | 135 | 26 | y (Hippocampal Sclerosis) | Right -Temporal |
| ID12 | 49 | 10 | 23 | 93 | 59 | y (space occupying amygdala) | Left-Temporal |
| ID13 | 92 | 7 | 19 | 100 | 49 | y (focal cortical dysplasia) | Right -Frontal |
| ID14 | 74 | 7 | 154 | 1002 | 36 | y (pilocytic astrocytoma) | Left-Parietal |
| ID15 | 61 | 3 | 52 | 184 | 23 | n | Left-Temporal |
| ID16 | 59 | 6 | 67 | 117 | 31 | y (Hippocampal Sclerosis) | Left-Temporal |

Supplementary Figure SF1: Figure depicts the robustness of results against varying the heuristically chosen bin size (bin size = 10 in the main manuscript). Plotted is the averaged Amplitude Entropy (AE) per frequency band before, during, and after seizure, as in Fig.3 of the main manuscript. The individual AE points are first averaged over seizures, then over patients (as illustrated in Fig.1), and finally, over time windows corresponding to before, during, and after a seizure. Note the change in scaling while showing the same triangular shape for different bin widths in (a) and (b).

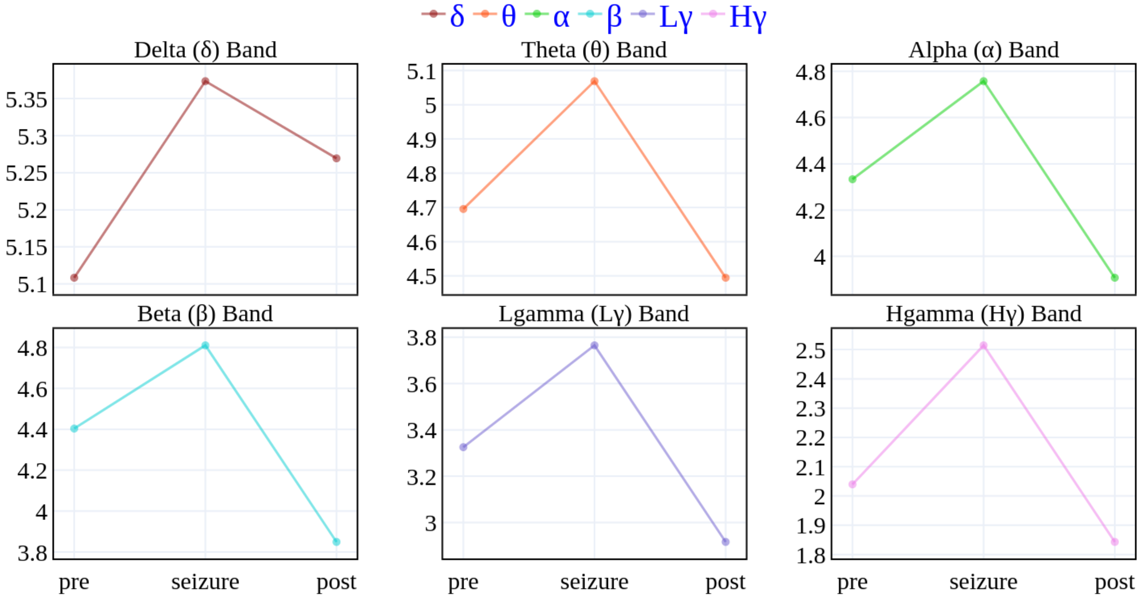


(a) Considered bin width 1 (a.u.)

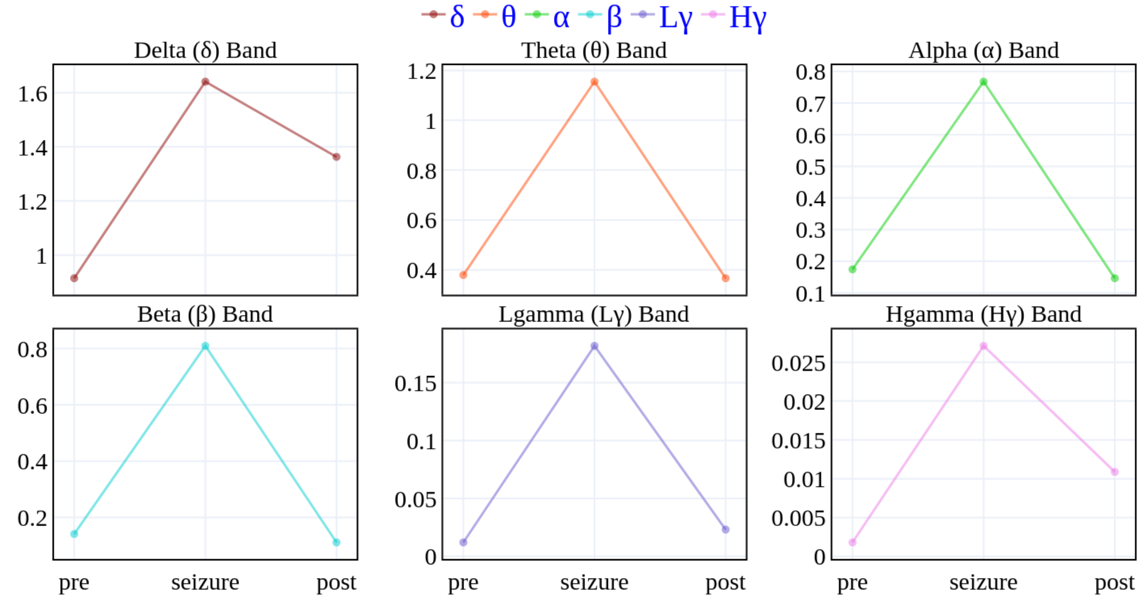


(b) Considered bin width 50 (a.u.)

Supplementary Figure SF2: This figure illustrates the distribution of the Analytic Amplitude (AA) across iEEG channels (panels **A** and **B**) and the evolution of Amplitude Entropy (AE) over time (panel **C**), grand-averaged across all seizures and patients, in this order, as in Fig.2 of the main manuscript, but for differently chosen bin width. Standard deviation averaged across patients in shades. Note the dfference in number of bins and mean AE for different bin widths in (a) and (b).

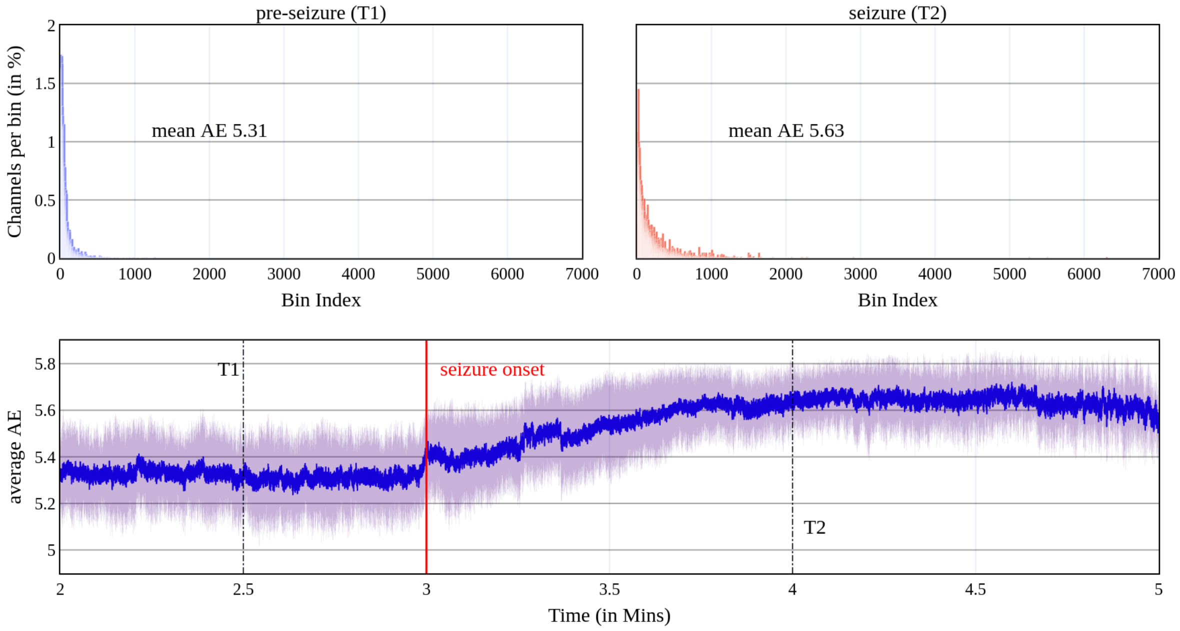


(a) Considered bin width 1 (a.u.)

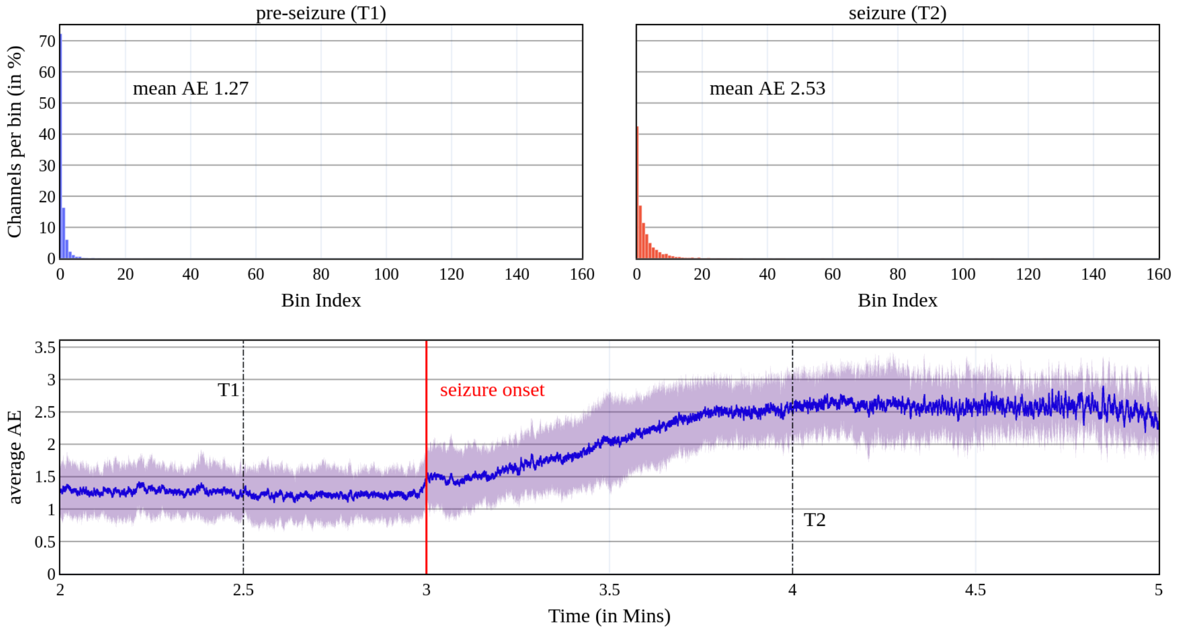


(b) Considered bin width 50 (a.u.)

Supplementary Figure SF3: Time-evolution of grand-averaged AE (averaged over both seizures and patients) as in Fig. SF7, when using the time normalization approach with regards to seizure duration as an alternative to the NaN padding procedure. The average shows a clear increase and plateau around 0.5 of the total seizure duration, in line with our speculation that average AE might increase during the progressive recruitment of brain areas during seizure propagation. See K. Schindler et al. (doi:10.1093/brain/awl304) for details about time normalization methodology.

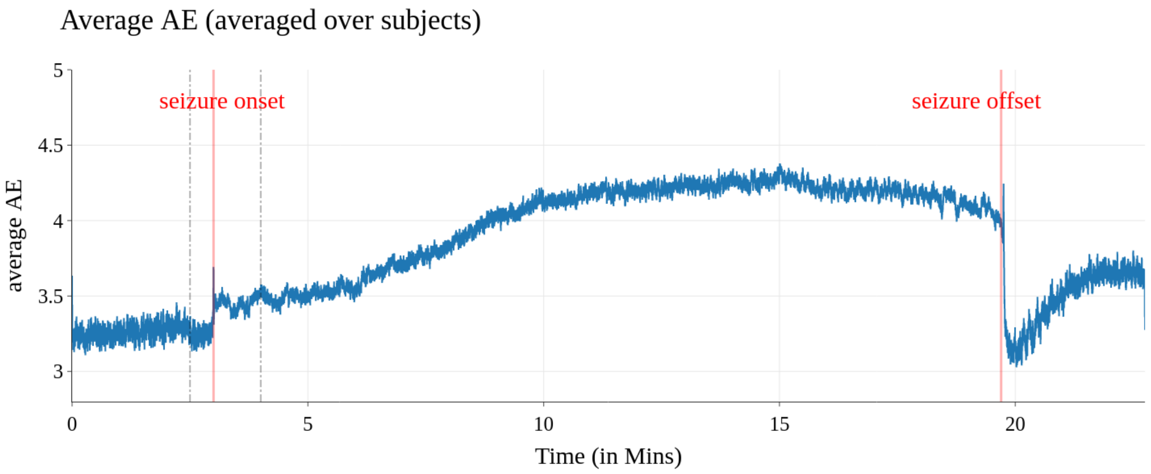

Supplementary Figure SF4: Time-evolution of Amplitude Entropy (AE); Each AE time-series is patient-resolved and averaged over seizures for each patient, as in Fig. SF6, when using the time normalization approach with regards to seizure duration as an alternative to the NaN padding procedure. See K. Schindler et al. (doi:10.1093/brain/awl304) for details about time normalization methodology.

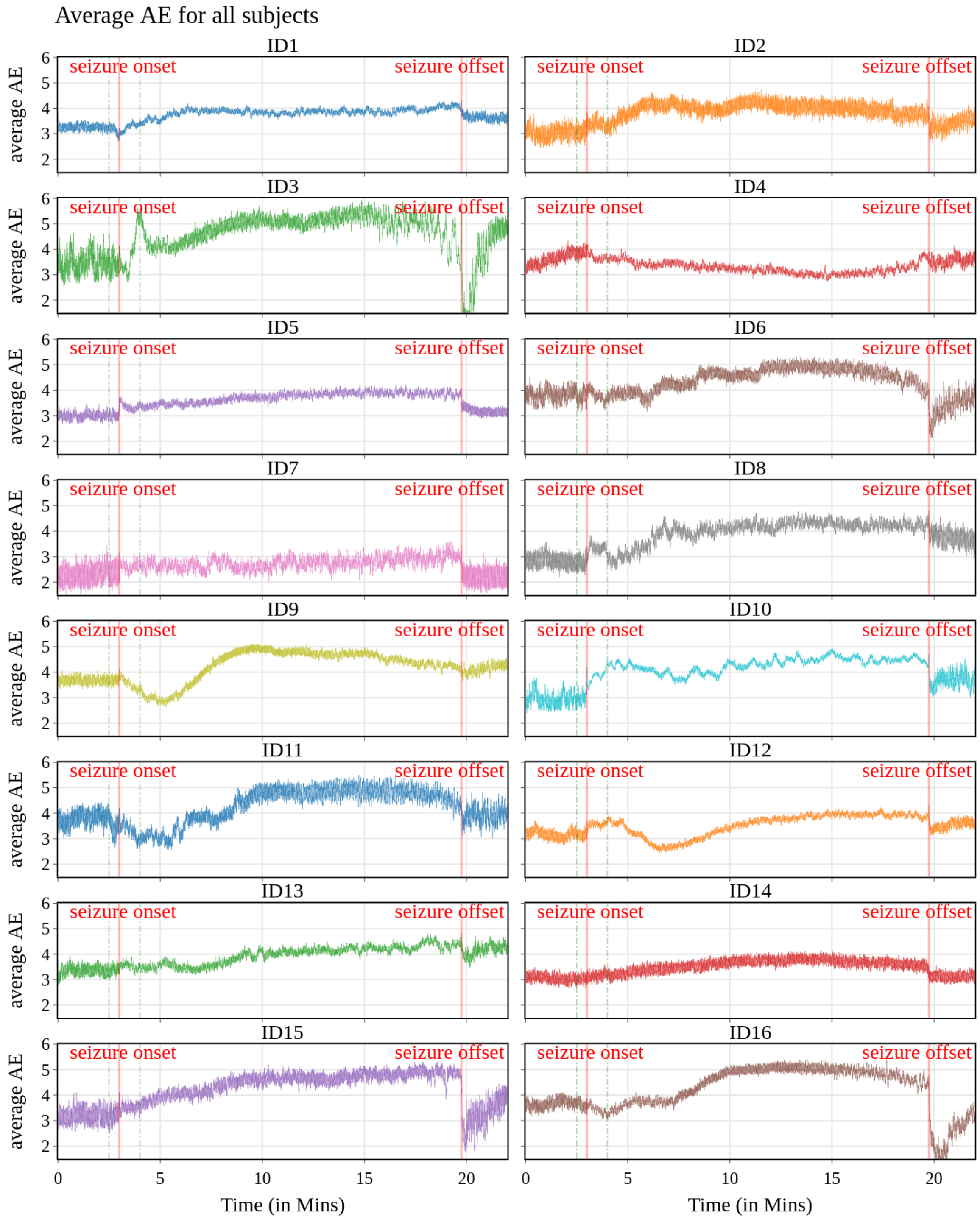

Supplementary Figure SF5: Distribution of the Analytic Amplitude (AA) of iEEG channels at T1 & T2 time-points as marked in Fig. SF6, averaged across seizures for each patients. Note for patient 7 and 10, due to short seizures, T2 outside seizure periods and thus not represented here.

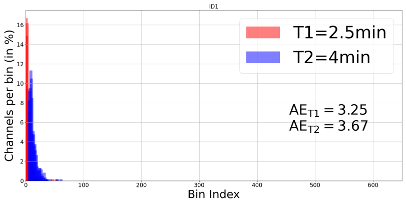

(a) Patient 1

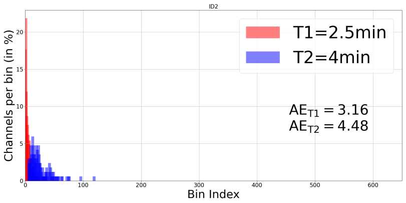

(b) Patient 2

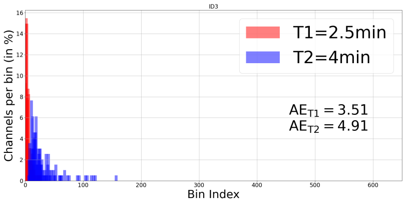

(c) Patient 3

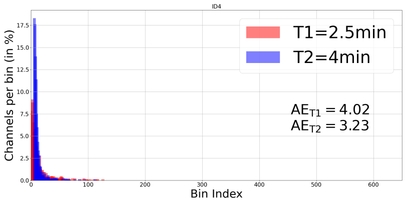

(d) Patient 4

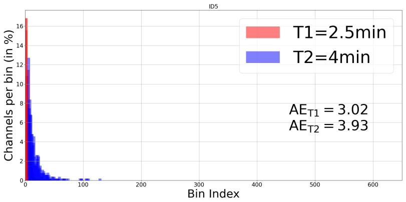

(e) Patient 5

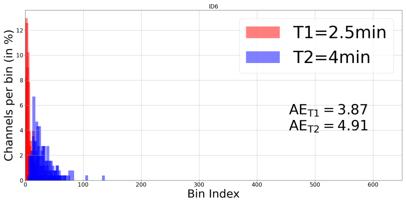

(f) Patient 6

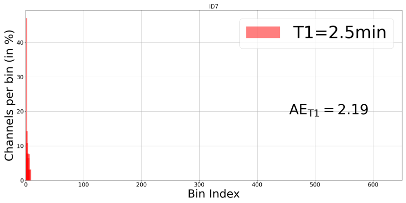

(g) Patient 7

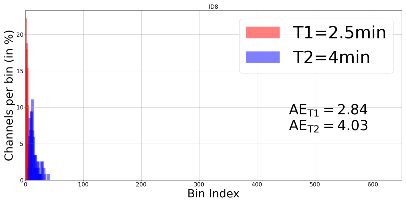

(h) Patient 8

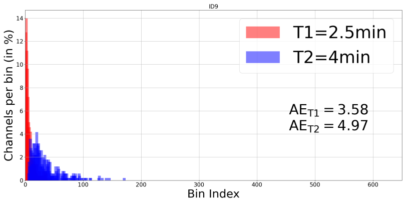

(i) Patient 9

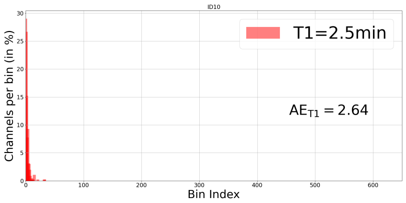

(j) Patient 10

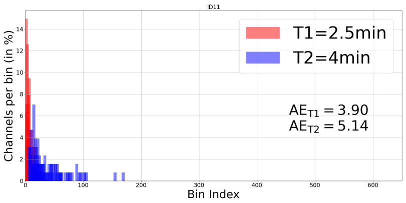

(k) Patient 11

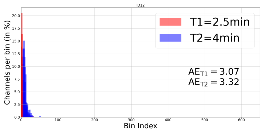

(l) Patient 12

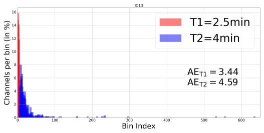

(m) Patient 13

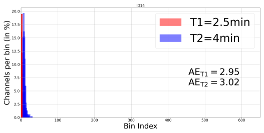

(n) Patient 14

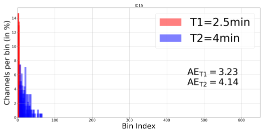

(o) Patient 15

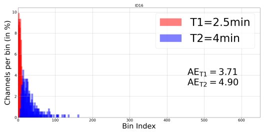

(p) Patient 16

Supplementary Figure SF6: Time-evolution of Amplitude Entropy (AE); AE time-series are averaged over seizures for each patient. Two time points namely $t = 2.5$ and $t = 4$ min are marked at *before* and *during seizure* (except ID 7 and 10) which corresponds to T1 & T2 in Fig.**2**(C) in the main manuscript. Note for patient 7 and 10, maximum seizure duration is 16 and 22 s, respectively (See table ST1). Due to such short seizures, in both cases, the values at T2 for ID 7 and 10 are padded with *NaN*s for averaging operation.

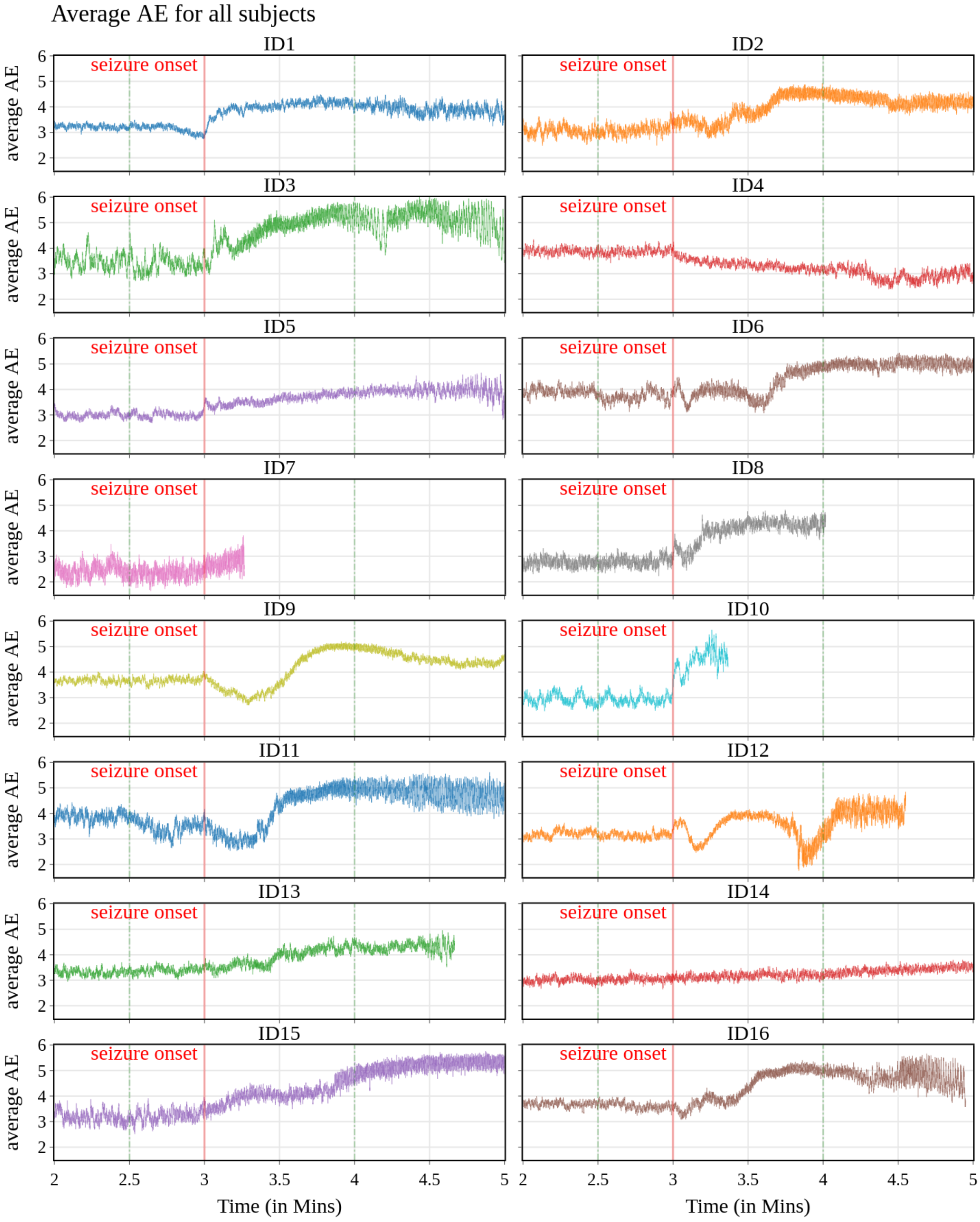

Supplementary Figure SF7: Time-evolution of averaged AE (averaged over both seizures and patients) with (**a**) including the largest seizure with time window of 1002 seconds ($\approx 17$ mins) (**b**) including upto the second largest seizure with a time window of 301 seconds ($\approx 5$ mins) in the seizure segment; both contains 3 min. of pre and post seizure data. Due to variable length of the seizures, (**a**) a maximum length seizure (of 1002 seconds) is chosen as the baseline time-series and all other seizures are stacked with a NaN padding to match the length of the baseline seizure. NaN mean was used for the averaging. The averaging is performed first seizure wise per patient and then across all the patients. Point to note here that the sharp drops that can be observed in averaged AE stems from the drop in number of samples in the seizure time-period. Since only one seizure is stretched for $1002$ seconds, most of the series in the seizure period is consisted of only one sample. This is the same AE series as displayed in Fig. **2**C of the main manuscript but with full length time series. Two time points is marked at before and during the seizure which corresponds to T1 & T2 in Fig. **2**C. (**b**) the averaging is performed on a slice of 301 secondsin the seizure window (with NaN padding) and 3 min. of pre. and post. data,separately and stacked for plotting.

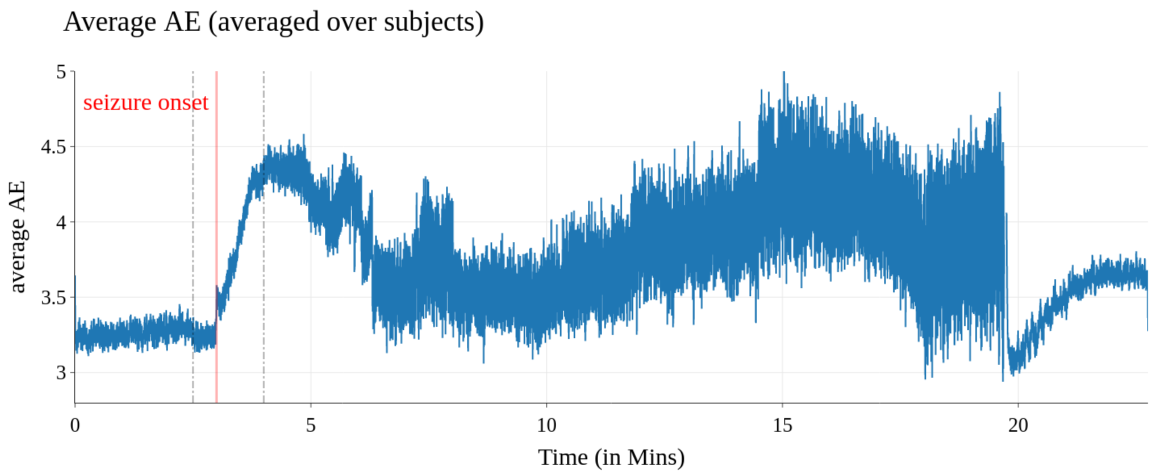


(a) Grand-averaged AE across all seizures.

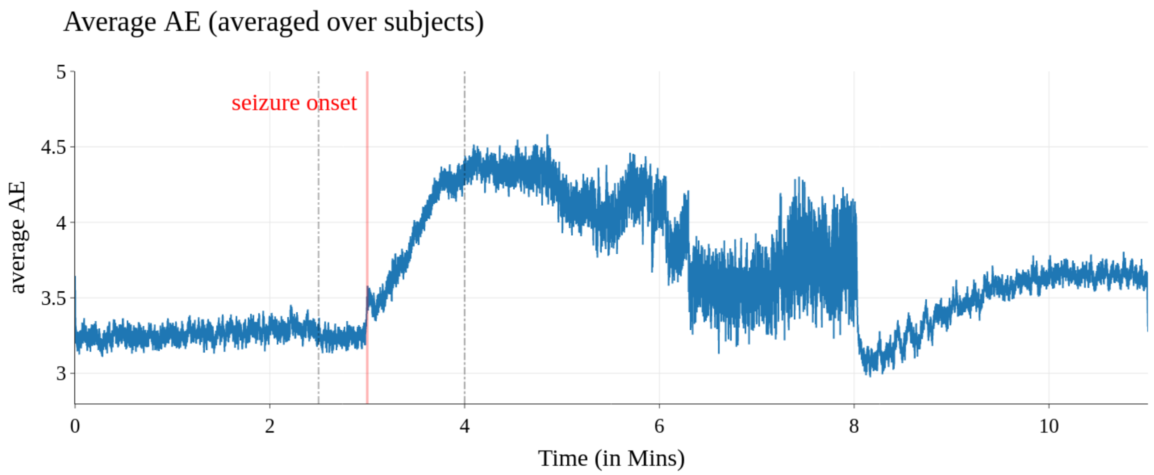


(b) Grand-averaged AE across all seizures excluding one seizure of patient ID14, that was exceptionally long (1002 s, see ST1)

Supplementary Figure SF8: This figure depicts the iEEG signals for **first seizures** for patients 1 to 6 as acquired from the SWEC data-set, showing relative variations between patients and seizures.

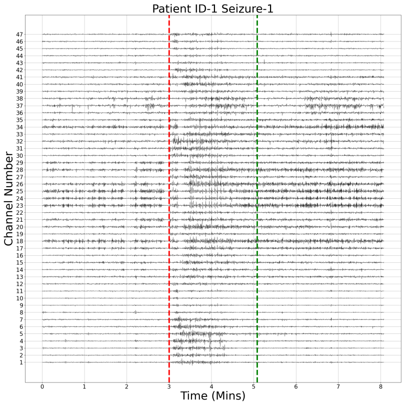


(a) Patient 1

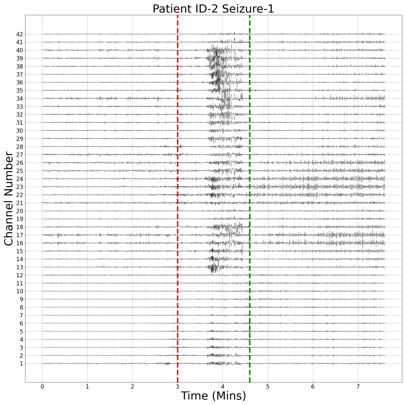


(b) Patient 2

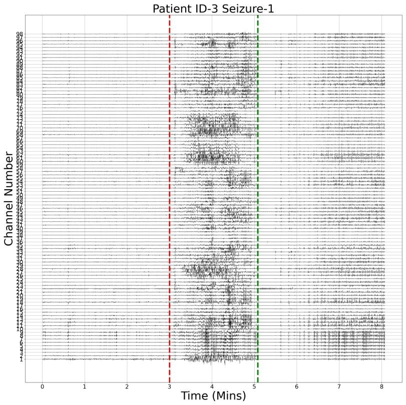


(c) Patient 3

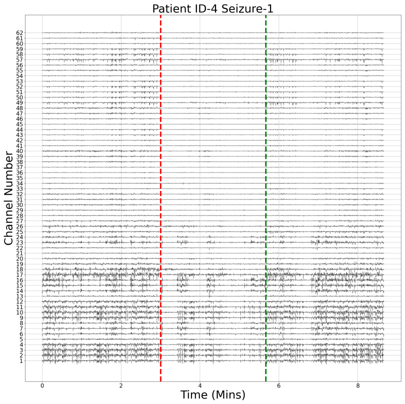


(d) Patient 4

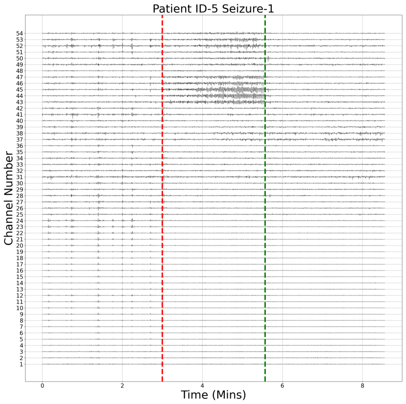


(e) Patient 5

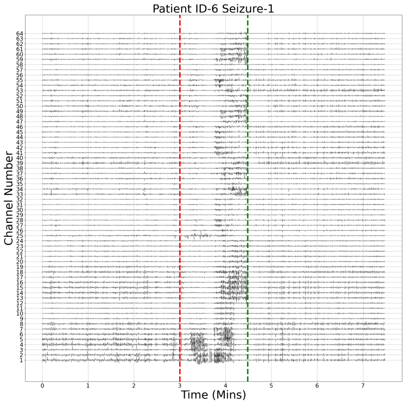


(f) Patient 6

Supplementary Figure SF9: This figure depicts the iEEG signals for **first seizures** for patients 7 to 12 as acquired from the SWEC data-set, showing relative variations between patients and seizures.

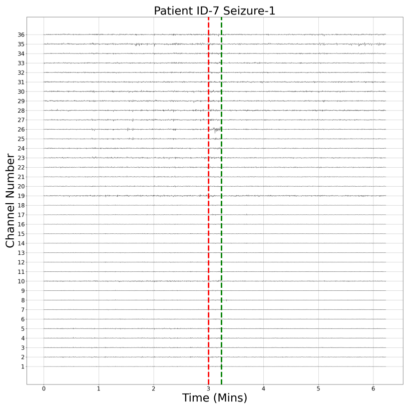


(a) Patient 7

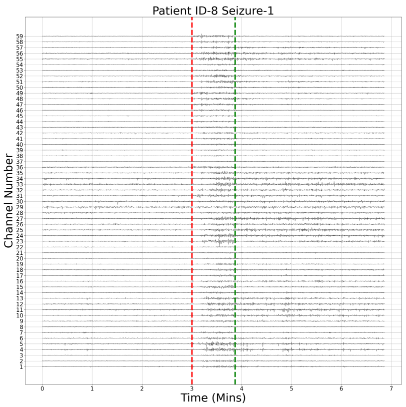


(b) Patient 8

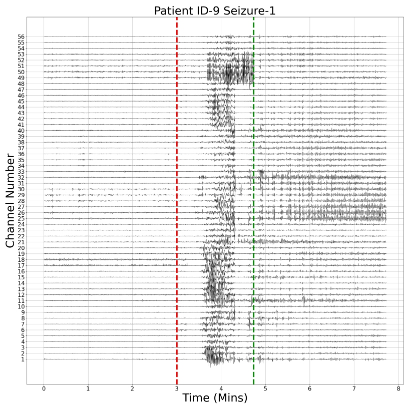


(c) Patient 9

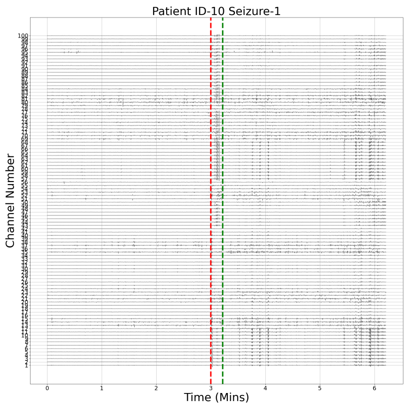


(d) Patient 10

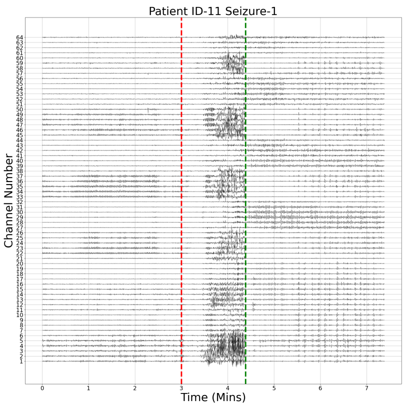


(e) Patient 11

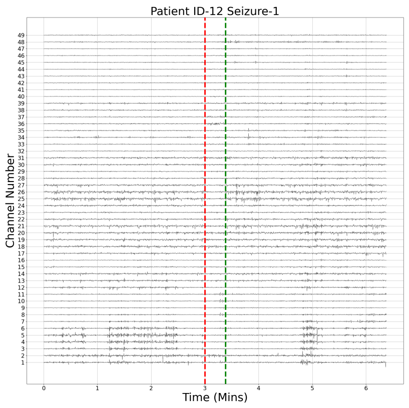


(f) Patient 12

Supplementary Figure SF10: This figure depicts the iEEG signals for **first seizures** for patients 13 to 16 as acquired from the SWEC data-set, showing relative variations between patients and seizures.

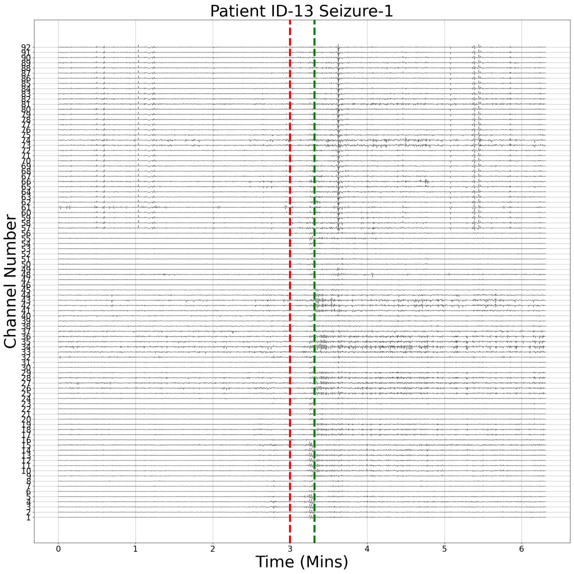


(a) Patient 13

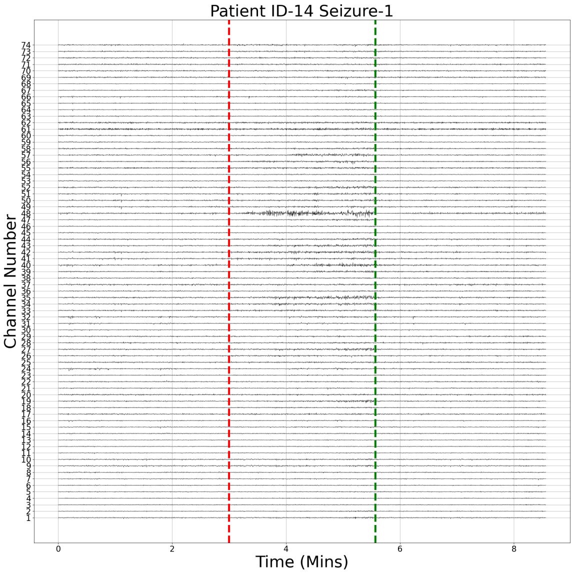


(b) Patient 14

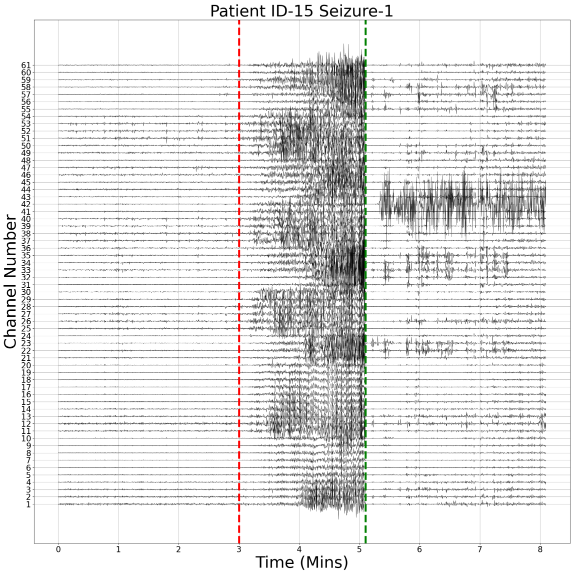


(c) Patient 15

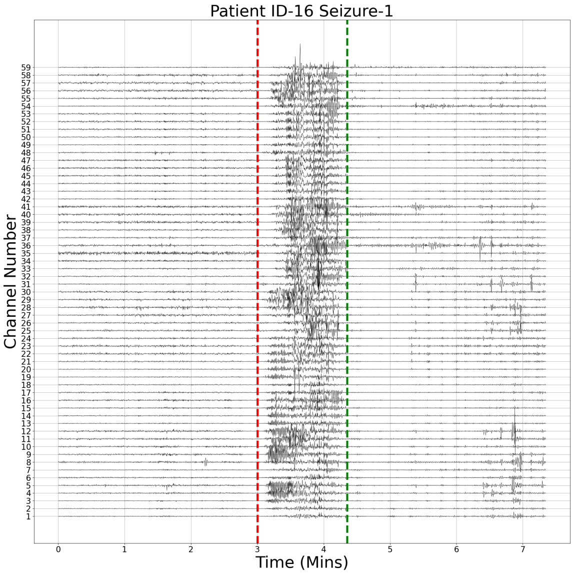


(d) Patient 16

Supplementary Figure SF11: This figure depicts the iEEG signals for **second seizures** for patients 1 to 6 as acquired from the SWEC data-set, showing relative variations between patients and seizures.

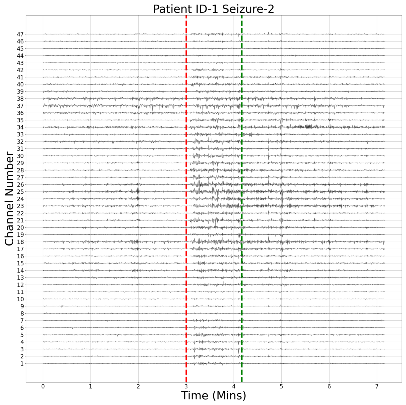


(a) Patient 1

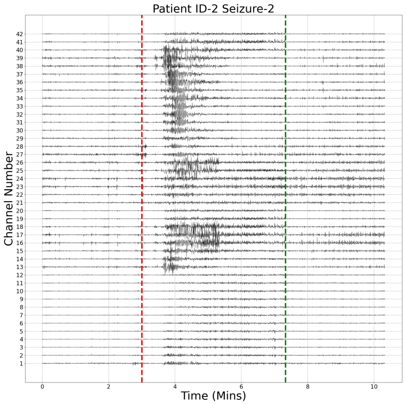


(b) Patient 2

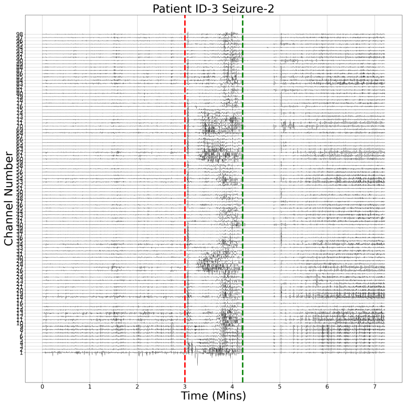


(c) Patient 3

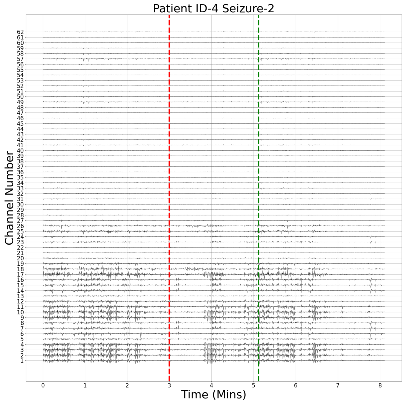


(d) Patient 4

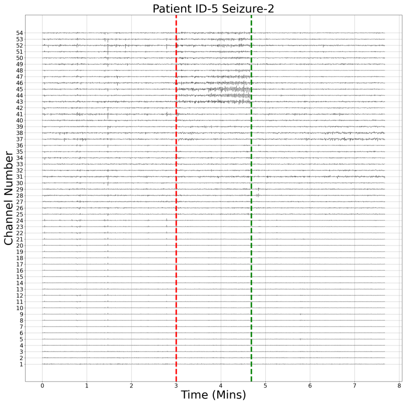


(e) Patient 5

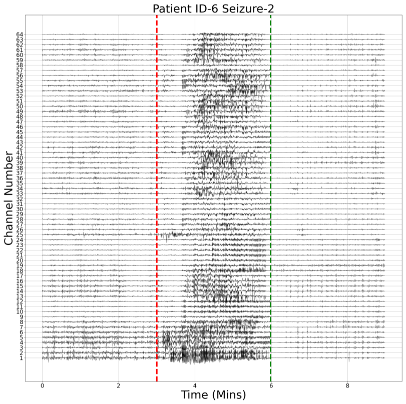


(f) Patient 6

Supplementary Figure SF12: This figure depicts the iEEG signals for **second seizures** for patients 7 to 12 as acquired from the SWEC data-set, showing relative variations between patients and seizures.

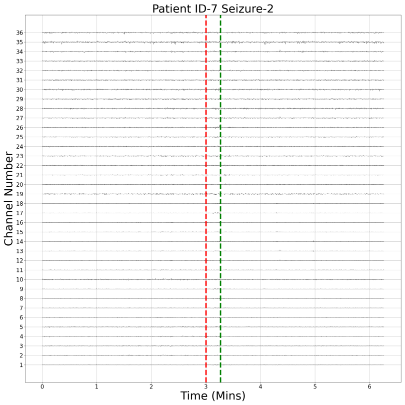


(a) Patient 7

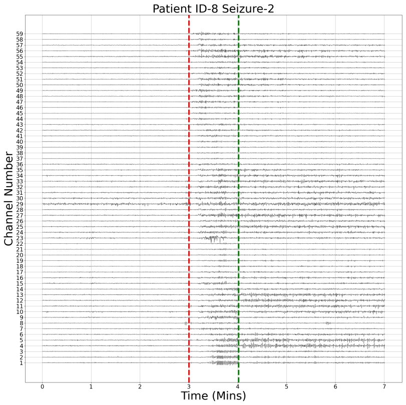


(b) Patient 8

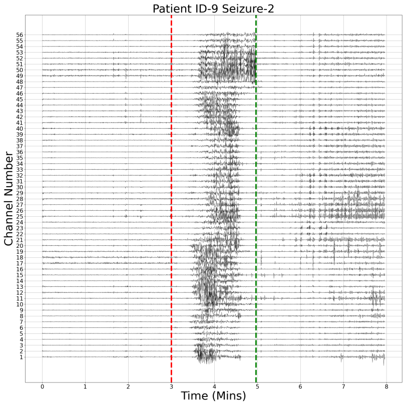


(c) Patient 9

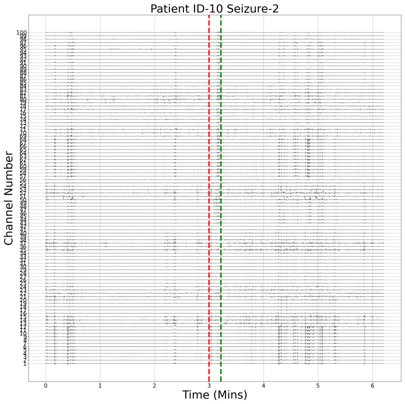


(d) Patient 10

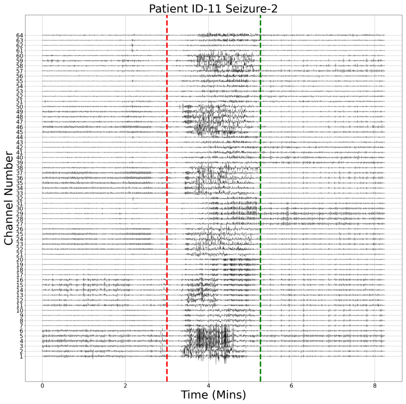


(e) Patient 11

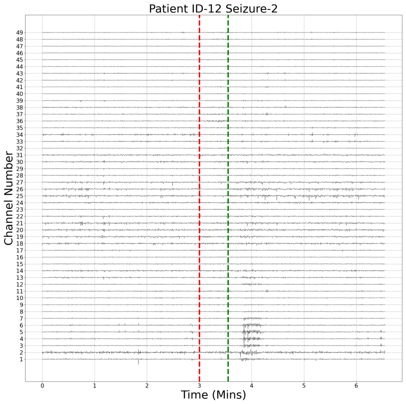


(f) Patient 12

Supplementary Figure SF13: This figure depicts the iEEG signals for **second seizures** for patients 13 to 16 as acquired from the SWEC data-set, showing relative variations between patients and seizures.

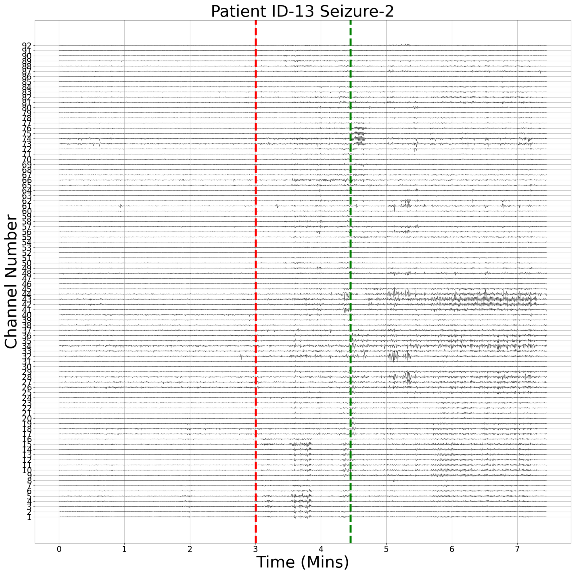


(a) Patient 13

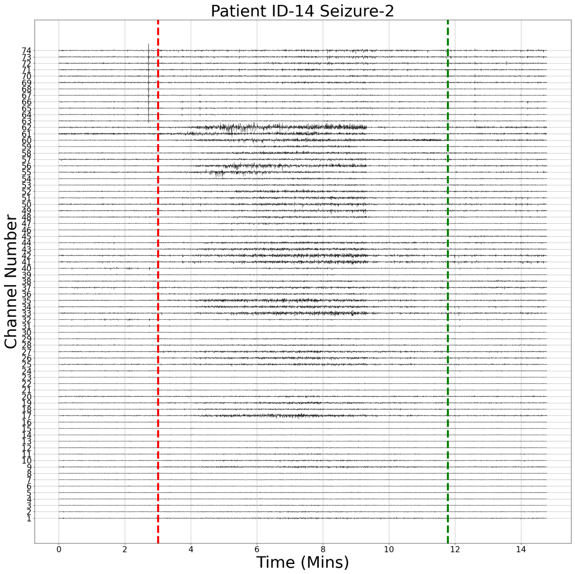


(b) Patient 14

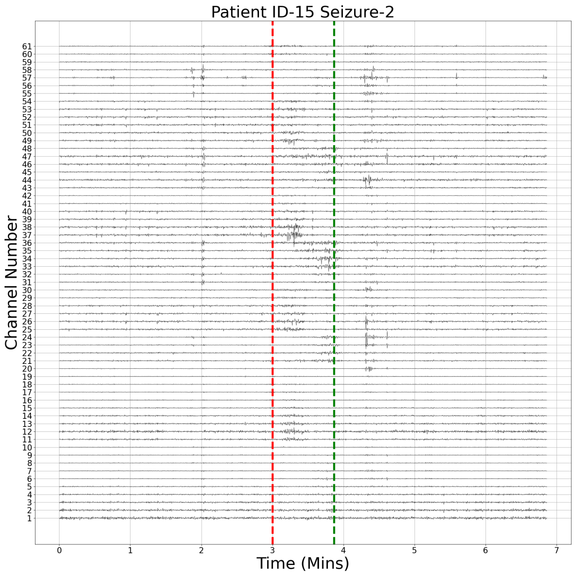


(c) Patient 15

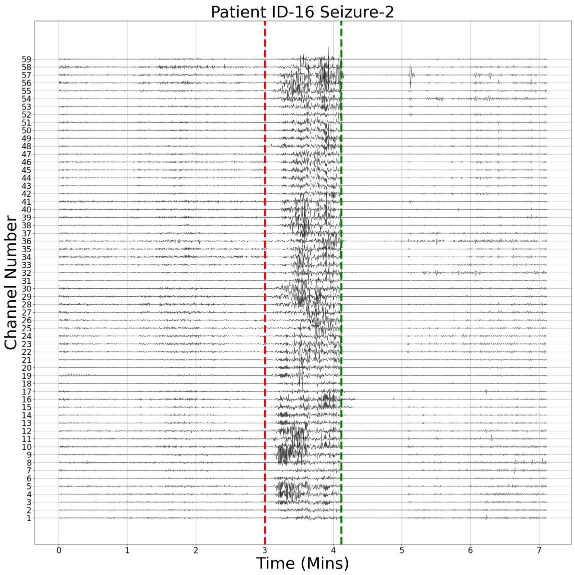


(d) Patient 16

Supplementary Figure SF14: This figure depicts the local peak in analytic amplitudes for different channels for Patient 2, Seizure 3.

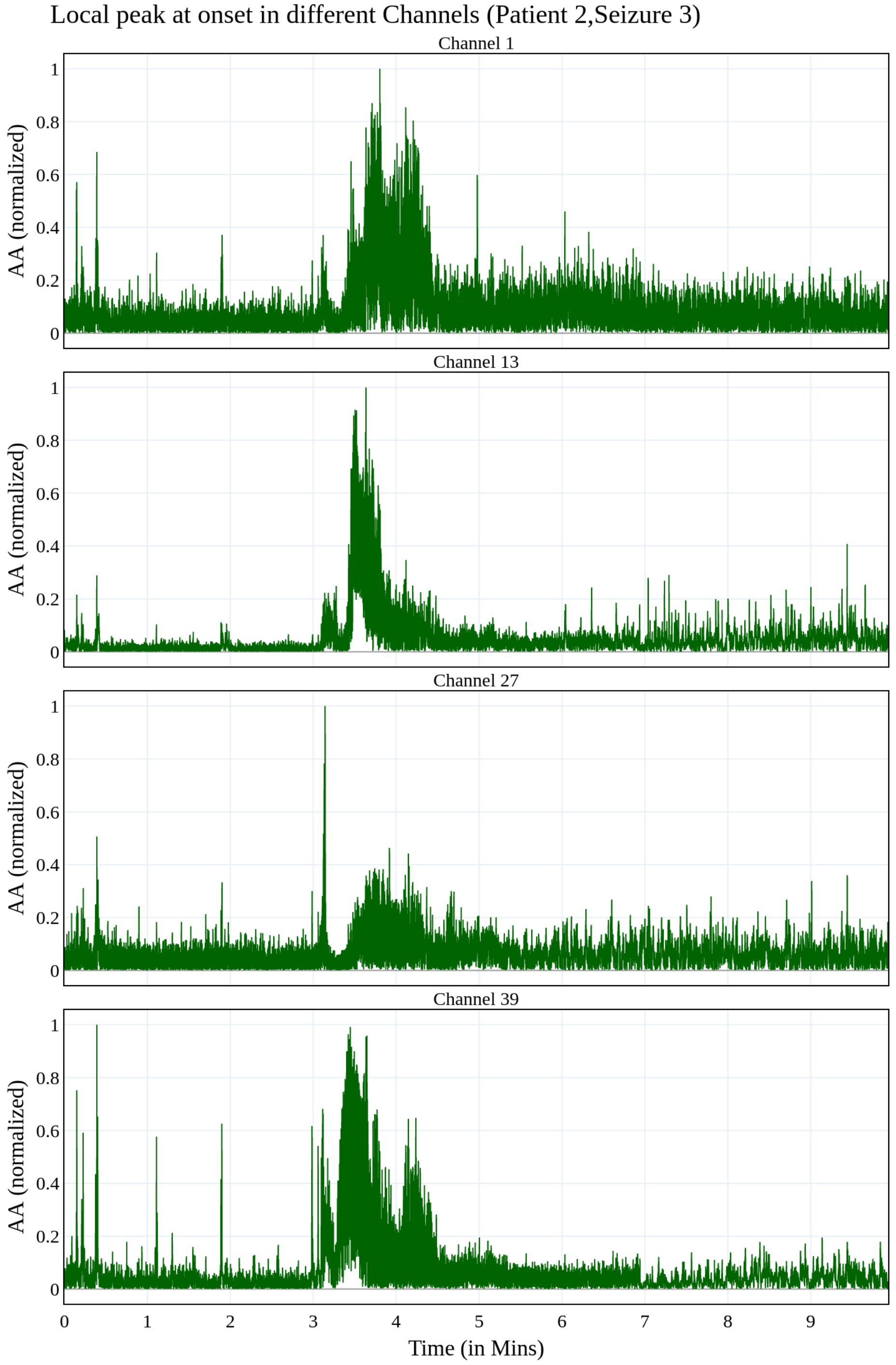

Supplementary Figure SF15: Frequency-resolved averaged AE (over seizures) before, during, and after a seizure per patient and frequency band. The individual AE points are averaged over seizures (as illustrated in Fig.1 in the main manuscript), and finally, over time windows corresponding to before, during, and after a seizure. The AE points exhibit a *significant* change during seizures compared to the time periods before or after the seizure.

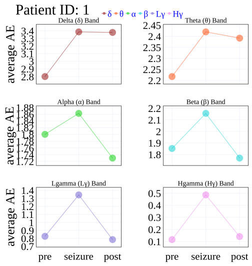

(a) Patient 1

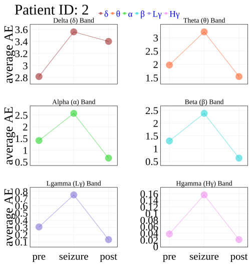

(b) Patient 2

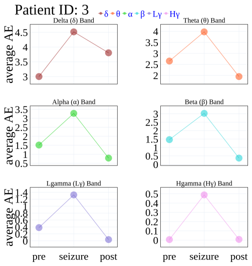

(c) Patient 3

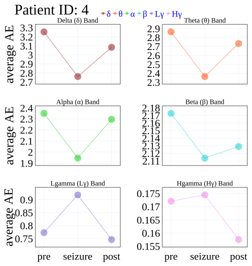

(d) Patient 4

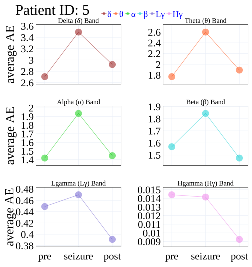

(e) Patient 5

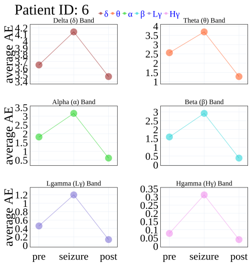

(f) Patient 6

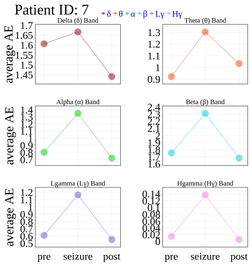

(g) Patient 7

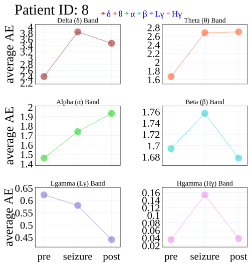

(h) Patient 8

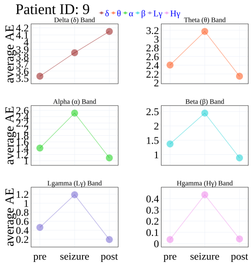

(i) Patient 9

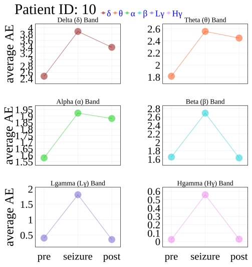

(j) Patient 10

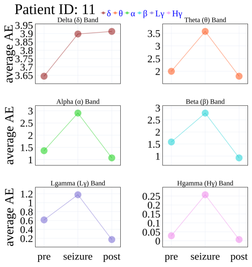

(k) Patient 11

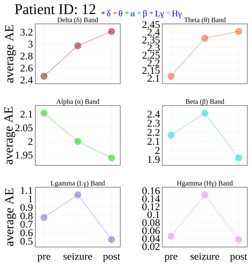

(l) Patient 12

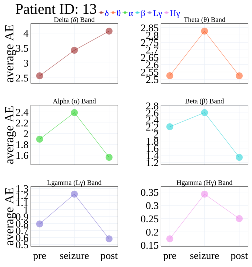

(m) Patient 13

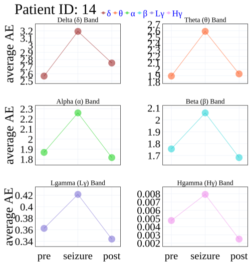

(n) Patient 14

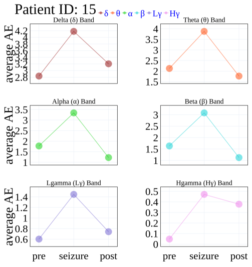

(o) Patient 15

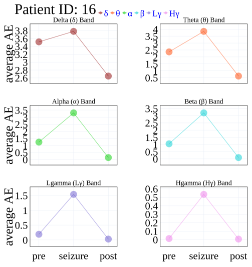

(p) Patient 16

Supplementary Figure SF16: Seizure effect on AE across patients presented in frequency separated and non-scaled fashion. The seizure effect is the difference between AE points corresponding to the seizure and before-seizure time periods (averaged). Patient-specific seizure effect distribution for AE calculated on filtered AA of iEEG channels for different frequency bands (color-coded, one dot per band per seizure).

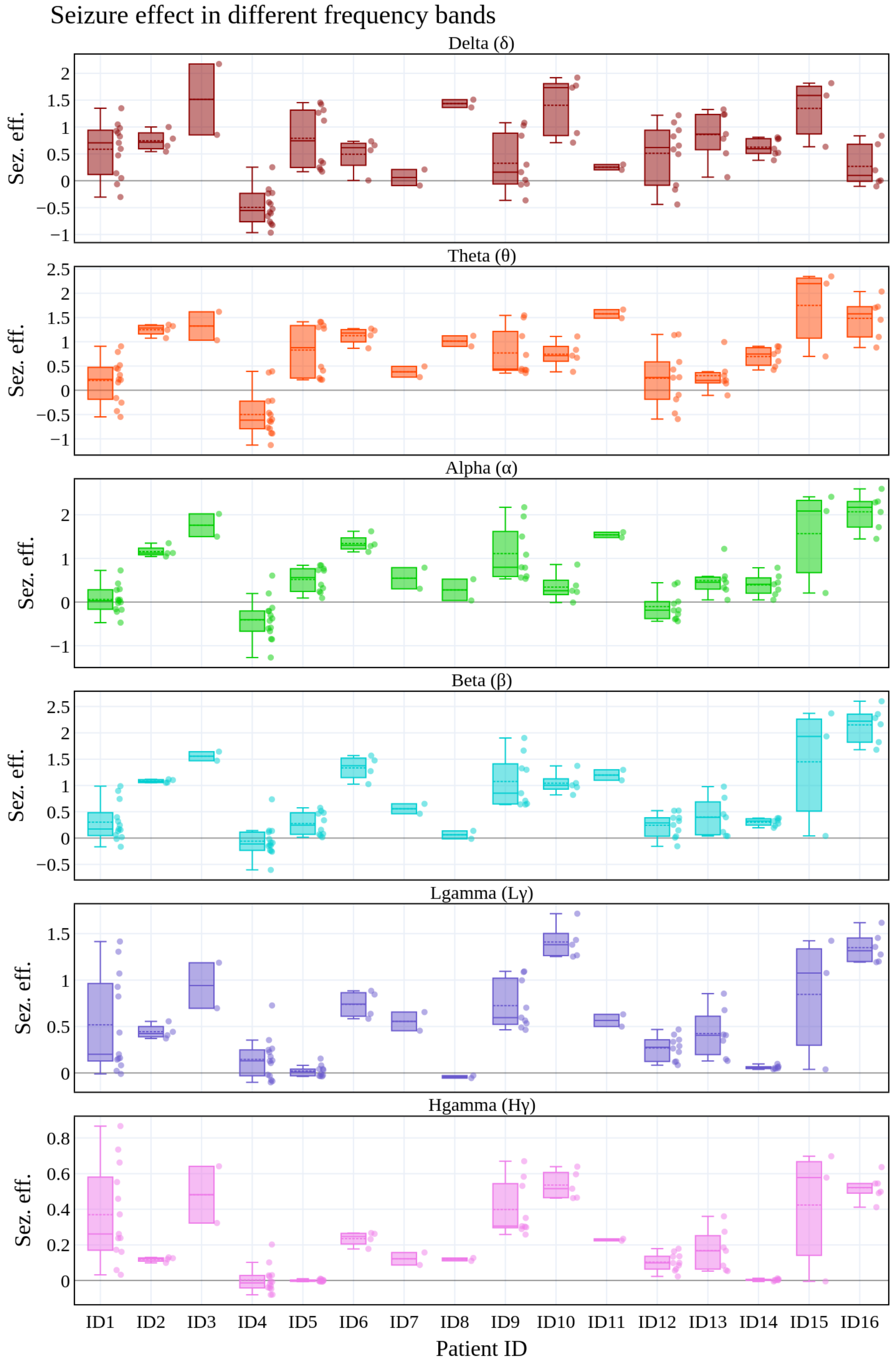
